# Neuronal-specific function of hTim8a in Complex IV assembly provides insight into the molecular mechanism underlying Mohr-Tranebjærg syndrome

**DOI:** 10.1101/725655

**Authors:** Yilin Kang, Alexander J. Anderson, David P. De Souza, Catherine S. Palmer, Kenji M. Fujihara, Tegan Stait, Ann E Frazier, Nicholas J. Clemons, Dedreia Tull, David R Thorburn, Malcolm J. McConville, Michael T. Ryan, David A. Stroud, Diana Stojanovski

## Abstract

Human Tim8a is a member of an intermembrane space chaperone network, known as the small TIM family, which transport hydrophobic membrane proteins through this compartment. Mutations in *TIMM8A* cause a neurodegenerative disease, Mohr-Tranebjærg syndrome (MTS), which is characterised by sensorineural hearing loss, dystonia and blindness. Nothing is known about the function of hTim8a in neuronal cells and consequently how lack of hTim8a leads to a neurodegenerative disease. We identified a novel cell-specific function of hTim8a in the assembly of Complex IV, which is mediated through a transient interaction with the copper chaperone COX17. Complex IV assembly defects in cells lacking hTim8a leads to oxidative stress and changes to key apoptotic regulators, including cytochrome c and Bax, which primes cells for cell death. Alleviation of oxidative stress using Vitamin E rescues cells from apoptotic vulnerability. We hypothesis that enhanced sensitivity of neuronal cells to apoptosis is the underlying mechanism of MTS.

## Introduction

Mitochondria are fundamental cellular organelles governing many metabolic processes including fatty acid oxidation, the Krebs cycle, oxidative phosphorylation (OXPHOS), and fatty acid ß-oxidation (Kasahara and Scorrano, 2014; McBride et al., 2006). Organelle dysfunction is associated with a broad spectrum of diseases, including diabetes, cardiovascular disease, neurodegenerative disease (Gorman et al., 2016; Nunnari and Suomalainen, 2012), and mitochondrial diseases that are genetic, often inherited disorders associated with energy generation defects (Frazier et al., 2017). The health and functionality of the mitochondrial network relies on the biogenesis of >1500 nuclear encoded proteins, which are imported into the organelle using sophisticated translocation machines (Chacinska et al., 2009; Endo and Yamano, 2009).

The small TIM proteins form an intermembrane space chaperone network that shield and transport hydrophobic membrane proteins (outer membrane beta-barrels and inner membrane metabolite carriers) as they passage the aqueous intermembrane space to membrane-localised translocation machines (Curran et al., 2002a; Curran et al., 2002b; Endres et al., 1999; Hoppins and Nargang, 2004; Wiedemann et al., 2004). Yeast have five small TIM chaperones, Tim8, Tim9, Tim10, Tim12 and Tim13 (Koehler et al., 1999b; Stojanovski et al., 2012), which assemble into hexameric complexes in the intermembrane space (Tim9-Tim10 or Tim8-Tim13) or at the inner membrane-located <u>T</u>ranslocase of the Inner <u>M</u>embrane 22 (TIM22) complex (Tim9-Tim10-Tim12) (Koehler, 2004; Koehler et al., 1999a; Koehler et al., 1998). Humans have six small TIM proteins that have been poorly characterised, these include: hTim8a, hTim8b, hTim9, hTim10a, hTim10b, and hTim13 (Bauer et al., 1999; Gentle et al., 2007). Why human mitochondria have two Tim8 isoforms is unclear, but hTim8a is predominately expressed in brain (Jin et al., 1999), suggesting tissue specific functions. In line with this, mutations in *TIMM8A* cause Mohr-Tranebjærg syndrome (MTS), an X-linked recessive neurodegenerative disorder characterised by progressive sensorineural hearing loss, dystonia, cortical blindness and dysphagia (Jin et al., 1996; Koehler et al., 1999a; Tranebjaerg et al., 1995).

We set out to investigate the function of hTim8a in human cells to gauge insight into the pathomechanism underlying MTS. To do this, we performed CRISPR/Cas9-genome editing of *TIMM8A* in: (i) SH-SY5Y cells, a human neuroblastoma cell line often used as an *in vitro* model of neuronal function; and (ii) the widely used HEK293 cells as a comparative control so we could dissect cell-specific functions (if any) of hTim8a. SH-SY5Y cells lacking hTim8a have defects in the assembly of Complex IV (cytochrome *c* oxidase), with associated loss of mitochondrial function and metabolic pathways that are specifically elevated in neurons, including the Aspartate/Malate shuttle. We show that hTim8a feeds into an interaction web of Complex IV chaperones and assembly factors, including the copper chaperones, COX17 and SCO1. Oxidative stress and changes to key apoptotic regulators, including cytochrome *c* and Bax, sensitises cells lacking hTim8a to intrinsic cell death and alleviation of oxidative stress using Vitamin E rescues cells from this apoptotic vulnerability. This data provides a molecular explanation for previously reported neuronal cell loss in MTS patients (Tranebjaerg et al., 2001) and suggests that early intervention with antioxidant could represent a treatment strategy for mitochondrial neuropathologies like Mohr-Tranebjærg syndrome.

## Results

### A novel and neuronal cell-specific function of hTim8a in Complex IV biogenesis

If hTim8a functions analogous to its yeast counterpart, Tim8a, we hypothesised that depletion or removal of the protein should have an impact on hydrophobic proteins of the inner and outer membranes (Hoppins and Nargang, 2004; Kang et al., 2016; Kang et al., 2017; Wiedemann et al., 2004). Therefore, we first set out to establish the substrate spectrum of the hTim8a-hTim13 complex, and utilised the hTim9-hTim10a complex as a comparative control. The *TIMM8A* and *TIMM9* genes were targeted using CRISPR/Cas9 in HEK293 to modify their expression. Genomic sequencing of the edited cells confirmed the presence of a 1-bp insertion in *TIMM8A* (**Supplemental Figure 1A**), while the *TIMM9* edited cells revealed two indel variants bearing a 1-bp deletion or 13-bp deletion, causing frame-shift mutations in the two alleles and new stop codons at 2 or 4 aa beyond the wildtype hTim9 stop codon (**Supplemental Figure 1B**). SDS-PAGE and immunoblot analysis on mitochondria isolated from these cells revealed the presence of a slower migrating hTim9 mutant protein that was significantly reduced at the steady-state level **(Figure 1A, left panel).** To the contrary mitochondria from the *TIMM8A* modified cells showed complete loss of the hTim8a protein (**Figure 1A, middle panel**). Given this, we refer to these cell lines as hTim9^MUT^ (MUT, mutant) and hTim8a^KO^ (KO, knockout). Given the neurological pathologies associated with mutation in *TIMM8A*, we also targeted *TIMM8A* in the neuronal-like SH-SY5Y cell line. We observed a mixture of modified and wild-type alleles in the isolated clone (**Supplemental Figure 1C**), however isolated mitochondria had no hTim8a visible by western blot (**Figure 1A, right panel**) and on mass spectrometric analyses (**Figure 1B, right panel**). It has been reported that the expression of hTim8a can be altered by skewed X-chromosome inactivation (Plenge et al., 1999), thus we hypothesise that the observed wild-type allele of *TIMM8A* is located on an inactive X-chromosome in these cells. As this cell line is not a complete knockout, we refer to it as hTim8a^MUT^.

**Figure 1.**
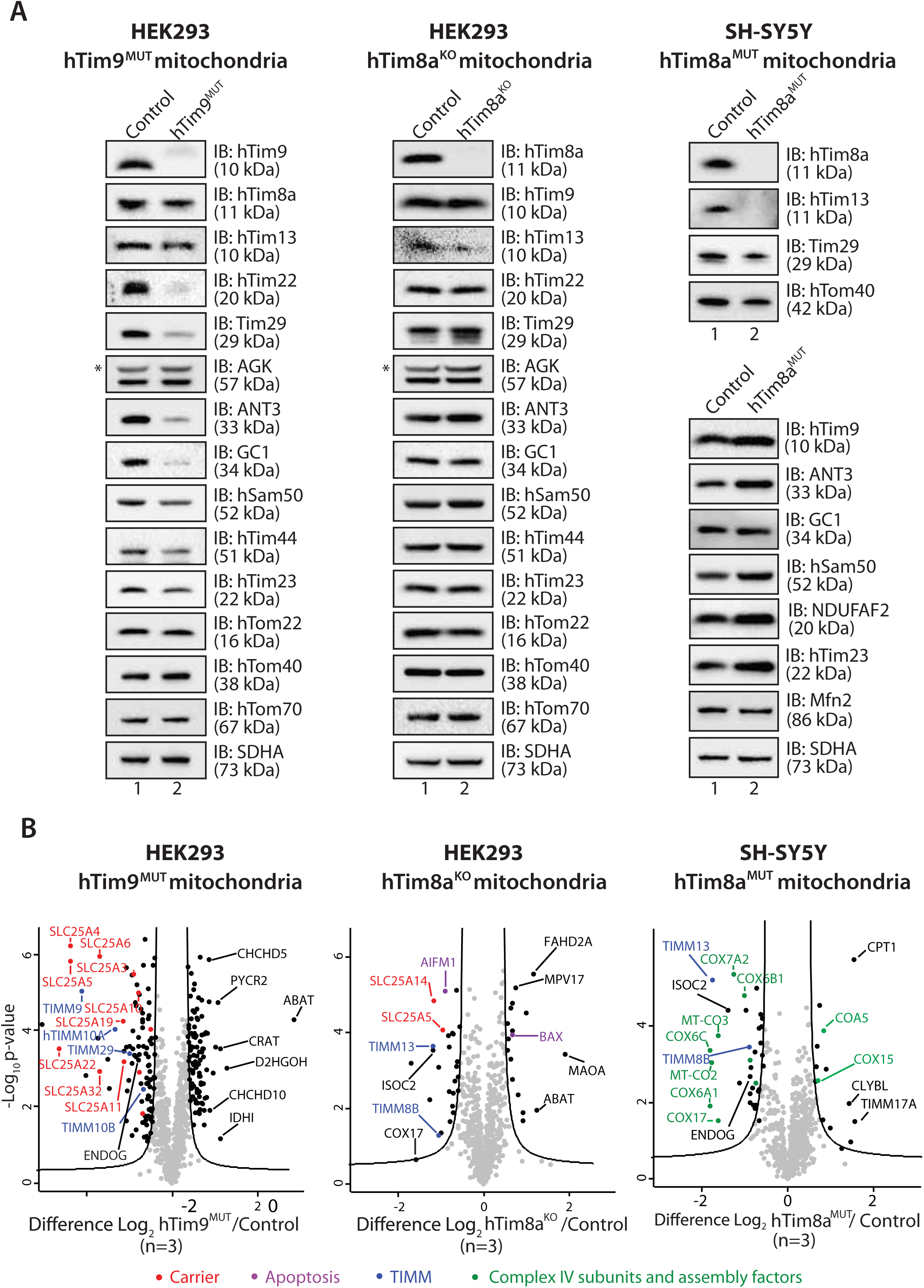
Cells lacking hTim8a have no defects in the TIM22 biogenesis pathway. (**A**) Mitochondria isolated from control, hTim9^MUT^ HEK293 cells (left panel), hTim8a^KO^ HEK293 cells (middle panel), or hTim8^MUT^ SH-SY5Y cells (right panel), were separated using SDS-PAGE for immunoblotting analyses with the indicated antibodies. **\*** on AGK panel indicates non-specific band of antibodies. (**B**) Mitochondria from (A) were subjected to label-free quantitative mass spectrometry. Volcano plots showing relative levels of proteins in knock-out cells compared to control cells. n=3 biological replicates. Significantly altered proteins are located outside the curved line (false discovery rate, FDR <5%): TIM22 complex subunits (blue), carrier proteins of the SLC25 family (red), apoptotic-related proteins (purple) and complex IV subunits and assembly factors (green). **See also Table 1.**

Having these tools in place, we addressed the functional implications of depleting hTim9 in HEK293 cells and hTim8a in both HEK293 and SH-SY5Y cells. Mitochondrial from hTim9^MUT^ cells resulted in a reduction in the steady state levels of TIM22 complex subunits (hTim22 and Tim29) and TIM22 substrate proteins (ANT3, GC1 and hTim23) (**Figure 1A, left panel)**, and displayed severe assembly defects of the TIM22 complex and the ANT3 dimer on BN-PAGE (**Supplemental Figure 2A**). To the contrary, lack of functional hTim8a in both HEK293 and SH-SY5Y cells had no obvious impact on the TIM22 complex or substrates when analysed by SDS-PAGE (**Figure 1A, middle and right panel**) or BN-PAGE (**Supplemental Figure 2B and 2C).** It has previously been suggested that that defects in hTim23 import and assembly into the TIM23 complex underlies MTS (Leuenberger et al., 1999; Paschen et al., 2000; Rothbauer et al., 2001). However, both hTim8^KO^ HEK293 and hTim8^MUT^ SH-SY5Y cells had no obvious impact on the TIM23 complex or Tim23 protein, suggesting an alternative pathomechanism. Taken together this data demonstrates that hTim9 is firmly embedded in the TIM22 complex biogenesis pathway, like its yeast counterpart, while hTim8a does not to influence the TIM22 complex or pathway and may have alternative function(s) within human mitochondria.

**Figure 2.**
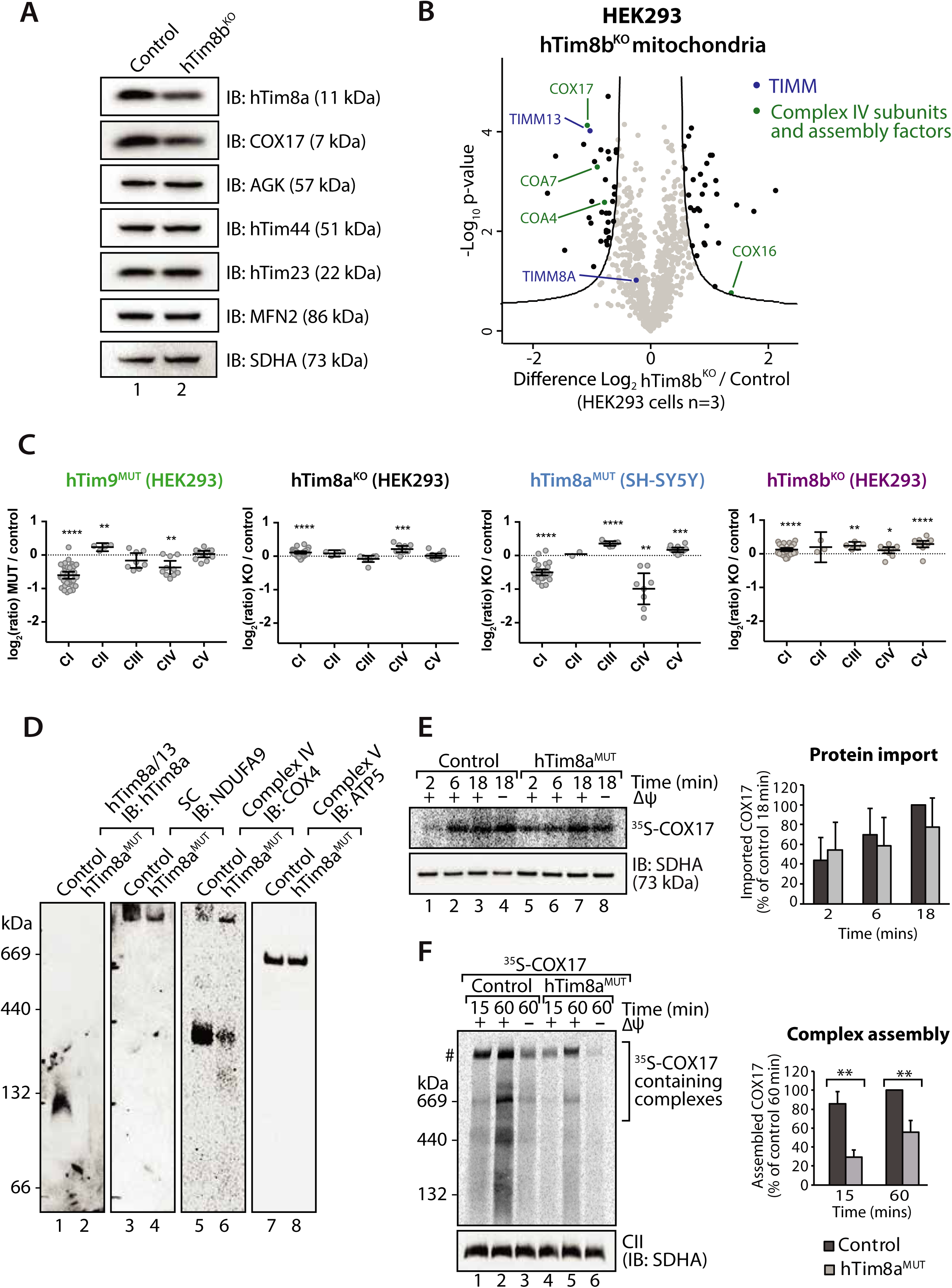
hTim8a and hTim8b show cell type specific function in Complex IV biogenesis. (**A and B**) Mitochondria from control and hTim8b^KO^ HEK293 cells were subjected to (A) SDS-PAGE/immunoblotting analyses or (B) label-free quantitative mass spectrometry. Volcano plot shows the relative levels of proteins in knock-out cells compared to control cells. n=3 biological replicates. Significantly altered proteins are located outside the curved line (false discovery rate, FDR <5%): TIM22 complex subunits (blue) and complex IV subunits and assembly factors (green). See also **Table 2**. (**C**) Relative abundance of respiratory chain complexes Complex I to V (subunits and assembly factors) in mitochondria isolated from control and hTim8a/8b/9-edited cells, detected using quantitative mass spectrometry in Figure 1B and 2B were quantified and tabulated as mean ± SEM (n=3). (**D**) Mitochondrial lysates from control and hTim8a^MUT^ SH-SY5Y cells were analysed using BN-PAGE and immunoblotting with the indicated antibodies. (**E and F**) ^35^S-labelled COX17 was incubated with mitochondria isolated from control and hTim8a^MUT^ cells in the presence or absence of membrane potential (*ΔΨ*) and treated with proteinase K prior to (**E**) TCA precipitation for SDS PAGE analysis, or (**F**) solubilised in 1% digitonin-containing buffer prior to BN-PAGE analysis, and autoradiography. The amount of (E) imported Cox17 (at 18 min) or (F) assembled COX17 (indicated by # on BN-PAGE at 60 min) were quantified and are represented as mean ± SD (n=3). **, p<0.01.

We performed label-free quantitative mass spectrometry on mitochondria isolated from hTim9^MUT^, hTim8^KO^ (HEK293) and hTim8^MUT^ (SH-SY5Y) to obtain a global view of the impact of their depletion on the mitochondrial proteome and mitochondrial health. hTim9^MUT^ mitochondria had reduced levels of hTim9 itself and its partner protein hTim10a, as small TIM are stable as hexameric units and not as monomers (Baker et al., 2012). In addition, TIM22 complex subunits, Tim29 and hTim10b, and numerous mitochondrial carrier proteins (SLC25 family), including ANT1 (*SLC25A4*) and the Phosphate Carrier (*SLC25A3*) were significantly reduced (**Figure 1B, left panel and Table 1**). Conversely, hTim8a^KO^ HEK293 mitochondria showed a decrease in the levels of: (i) hTim13 and hTim8b; (ii) the Apoptosis Inducing Factor, AIF; (iii) the Complex IV assembly factor, COX17; and (iv) two proteins belonging to the SLC25 family, SLC25A5 (ANT2) and SLC25A14, a previously described mitochondrial uncoupling protein (UCP5), also known as the brain mitochondrial carrier protein 1 (BMCP1) (Kim-Han et al., 2001; Sanchis et al., 1998) (**Figure 1B, middle panel and Table 1**). Mitochondria from hTim8a^KO^ cells also displayed upregulated levels of the pro-apoptotic BCL-2 family member, Bax. The changes to AIF and Bax were not evident in hTim9^MUT^ mitochondria, suggesting a specific cellular response due to the lack of hTim8a and a potential increased susceptibility to apoptosis. Looking at the proteomic changes in hTim8a^MUT^ SH-SY5Y cells (**Figure 1B, right plot and Table 1**), we noted only two substrates were commonly reduced in both HEK293 and SH-SY5Y cells lacking hTim8a: (i) ISOC2, and (ii) COX17 (**Figure 1B**). COX17 is a copper chaperone that delivers copper to Complex IV subunits, a crucial process for subsequent Complex IV assembly (Oswald et al., 2009). In line with this, hTim8a^MUT^ cells displayed a striking change to both mitochondrial and nuclear encoded Complex IV subunits and assembly factors (**Figure 1B, right panel**). Specifically, reduced levels of mitochondrially-encoded COX2 (*MT-CO2*) and COX3 (*MT-CO3*) and nuclear-encoded COX6A1, COX6B1, COX6C and COX7A2 were observed, while two putative assembly factors COA5 and COX15, were upregulated. This data suggested a cell specific function of hTim8a in neuronal-like cells in Complex IV biology. Given this, we set out to establish if the hTim8b isoform could be performing this role in HEK293 cells and indeed, hTim8b^KO^ HEK293 cells (**Supplemental Figure 1D**) showed reduced COX17 levels (**Figure 2A and 2B and Table 2**), in addition to Complex IV assembly factors COA4, COA7 and COX16 (**Figure 2B and Table 2**). A quantitative analysis of the proteomic data from both cell line models (**Figure 2C**) highlights a role for hTim9 (in HEK293 cells) in the biogenesis of respiratory chain complexes, in particular Complex I. Although hTim8a seems to have little impact on respiratory chain subunits and complexes in HEK293 cells, it has a prominent impact on Complex IV biogenesis in SH-SY5Y cells, highlighting a cell type-specific function of hTim8a in Complex IV biogenesis in neuroblastoma cells. To the best of our knowledge this is the first report of a cell specific function of a protein belonging to the mitochondrial protein import network.

### hTim8a transiently interacts with COX17 under high respiratory demand to promote Complex IV biogenesis/maturation in neuroblastoma cells

The common impact on COX17 in both HEK293 and SH-SY5Y lacking hTim8a, suggested a COX17-dependent process was being impacted in the absence of hTim8a. Mitochondria isolated from control and hTim8a^MUT^ SH-SY5Y cells were analysed by BN-PAGE and confirmed the lack of hTim8b-hTim13 complex, reduced levels of Complex I-containing supercomplex (probed with NDUFA9), and reduced levels of Complex IV (immunoblotting with COX4 antibodies) and preferential interaction of COX4 with supercomplexes in hTim8a^MUT^ mitochondria (**Figure 2D**). We investigated if hTim8a is required for the import and/or assembly of COX17 using *in vitro* import analysis. ^35^S-COX17 import into mitochondria isolated from hTim8a^MUT^ and control SH-SY5Y cells showed normal kinetics by SDS-PAGE (**Figure 2E**), which was expected given that COX17 is an established Mia40 substrate (Banci et al., 2010). However, analysis of invitro import reactions by BN-PAGE to monitor assembly if ^35^S-COX17 did suggest that COX17 assembly was severely perturbed in the absence of hTim8a (**Figure 2F;** ∼40% less COX17 assembly compared to control).

Previous reports have indicated that growth of cells on galactose enhances their oxidative capacity and that these cells exhibit increases in Complex IV activity and expression (Aguer et al., 2011; Rossignol et al., 2004). We cultured control and hTim8a^MUT^ SH-SY5Y cells in glucose or galactose-containing media (**Figure 2A, left schematic**) and analysed mitochondria isolated form these cells by SDS-PAGE and immunoblotting. Both control and hTim8a^MUT^ cells increased COX4 protein expression (Complex IV subunit) (**Figure 3A**) when cultured in galactose. However, this trend was also apparent for COX17 in hTim8a^MUT^ cells (**Figure 3A compare lanes 3 and 4**), suggesting that COX17 biogenesis is perturbed in cells lacking hTim8a. We set out to investigate if hTim8a was interacting with COX17 to accommodate a COX17-dependent process in mitochondria. hTim8a^FLAG^ was re-expressed in hTim8a^MUT^ SH-SY5Y cells and these cells were cultured in galactose prior to mitochondrial isolation, cross-linking with dithiobis-succinimidyl propionate (DSP) and immunoprecipitation using FLAG antibodies (**Figure 3B**). Using this approach, we captured an interaction between hTim8a^FLAG^ and COX17 (**Figure 3B)**. Label free quantitative mass spectrometry analyses of the hTim8a^FLAG^ cross link fractions revealed enrichment of multiple Complex IV metallochaperones (SCO1), assembly factors (CMC1, COA4, COA6, COA7 and COX6B1) and core subunit (COX4l1) (**Figure 3C and Table 3**), which are all involved in the early stage of Complex IV assembly. Strikingly, we detected a significant enrichment of SCO1 (**Figure 3C and Table 3**), a metallochaperone that is known to directly transfer copper from COX17 to the core component, COX2 of Complex IV (Leary et al., 2009). Collectively, these observations support a novel role of hTim8a in the maturation of Complex IV through an interaction with COX17 in SH-SY5Y cells.

**Figure 3.**
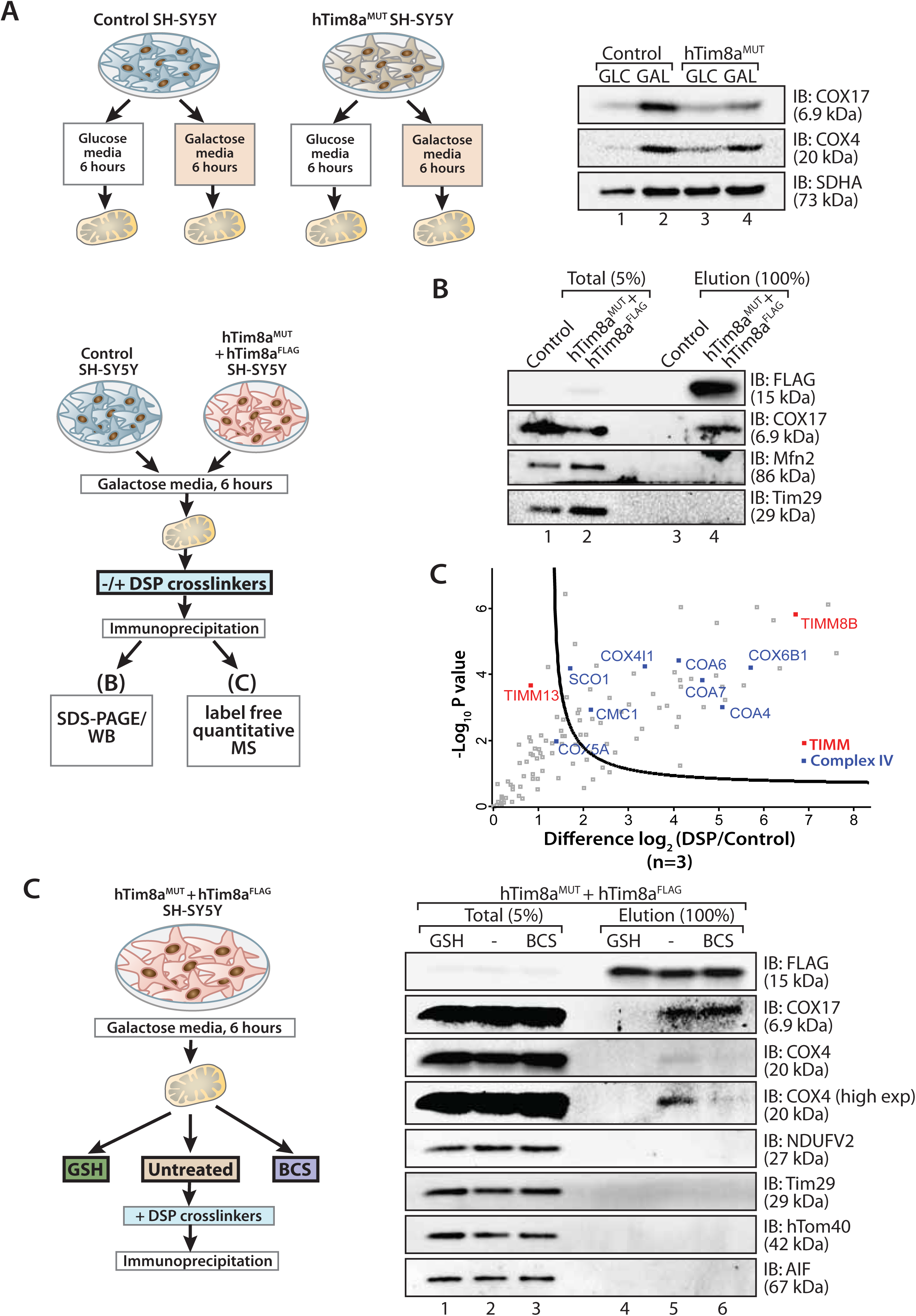
hTim8a functions in the early stage of assembly of Complex IV in SH-SY5Y cells. (**A**) Control or hTim8a^MUT^ SH-SY5Y cells were cultured in glucose-rich (GLC) or galactose (GAL) media for 6 hours prior to mitochondrial isolation for SDS-PAGE and immunoblotting analyses. (**B and C**) Mitochondria were isolated from control or hTim8a^MUT^ cells re-expressing hTim8a^FLAG^ (hTim8a^MUT^ + hTim8a^FLAG^) SH-SY5Y cells were incubated in doxycycline-containing media for 20 hours to induce hTim8a^FLAG^ expression. Cells were shifted to galactose media for 6 hours. Mitochondria were isolated from these cells and treated for crosslinking with dithiobis-succinimidyl propionate (DSP) and downstream immunoprecipitation prior to analyses using (B) SDS-PAGE and immunoblotting or (C) label free quantitative mass spectrometry. (**D**) hTim8a^MUT^ + hTim8a^FLAG^ mitochondria (protein expression induced with doxycycline for 20 hours followed by 6 hours incubation in galactose media) were either left untreated, subjected to bathocuproine disulfonate (BCS) chelation or reduced using glutathione (GSH) prior to DSP crosslinking and immunoprecipitation. Total and eluate fractions were separated using SDS-PAGE and analysed using immunoblotting.

Given the protein interaction profile of hTim8a in SH-SY5Y cells we questioned if it was functioning in the copper transfer pathways mediated by COX17 and SCO1. However, both COX17 and hTim8a are also MIA substrates that contain cysteine motifs that form disulfide bonds in the mature proteins (Stojanovski et al., 2012). To interrogate the nature of the hTim8a^FLAG^-COX17 interaction and whether it is reliant on copper or disulfide bonds, we pre-treated isolated mitochondria with either: (i) the copper chelator, bathocuproine disulfonate (BCS), or (ii) reduced glutathione (GSH) to reduce the disulfide bonds prior to crosslinking/immunoprecipitation (**Figure 3C)**. This revealed that the transient interaction between hTim8a and COX17 was not influenced by BCS suggesting that copper is not critical for the protein association. However, there was a pronounced reduction in the interaction of COX17 with COX4 upon copper chelation (**Figure 3C, compare lanes 5 and 6**), showing the validity of the approach. Importantly, pre-incubation of mitochondria with GSH significantly compromised the interaction of hTim8a^FLAG^ with COX17 (**Figure 3C, compare lanes 4 and 5**), indicating that the oxidation status of the cysteine residues in these proteins is crucial to maintain their association within mitochondria. Together, these data show that hTim8a interacts with COX17 in a redox sensitive-manner, and disruption of the hTim8a-COX17 interaction perturbs the downstream Complex IV assembly. The presence of COX4 in the control eluate fraction together with COX17 (**Figure 3C, lane 5**) suggests that hTim8a is binding to the COX17 pool participating in copper delivery to Complex IV (**Figure 3C**) and loss of this interaction perturbs maturation of Complex IV.

### Loss of hTim8a induces mitochondrial dysfunction and changes to mitochondrial metabolism

Complex IV dysfunction is linked to increased mitochondrial ROS and cellular toxicity (Srinivasan and Avadhani, 2012), thus we assessed mitochondrial membrane potential and oxidative stress in SH-SY5Y lacking hTim8a. We used hTim8^KO^ HEK293 cells as a control to compare the levels of mitochondrial dysfunction, if any. Incubation of cells with the membrane potential indicator, TMRM, indicated a compromised membrane potential in both SH-SY5Y and HEK293 cells lacking hTim8a, which was slightly more severe in the neuronal-like cells (**Figure 4A**). This was not due to perturbed mitochondrial biogenesis as citrate synthase activity, which is used as an indicator of mitochondrial content, showed mitochondrial volume was greater in hTim8a^KO^ cells (**Supplemental Figure 3A**). To assess oxidative stress, controls, hTim8a^KO^ and hTim8^MUT^ cells were either left untreated or pre-treated with the oxidative-stress inducing agent, menadione prior to H_2_O_2_ ROS level measurement. ROS levels were significantly elevated compared to controls in both SH-SY5Y and HEK293 cells (**Figure 4B**) confirming enhanced oxidative stress in the absence of hTim8a. Intriguingly, in spite of this mitochondrial dysfunction, both SH-SY5Y and HEK293 cells lacking hTim8a grew at a rate comparable to wild-type cells (**Figure 4C**).

**Figure 4.**
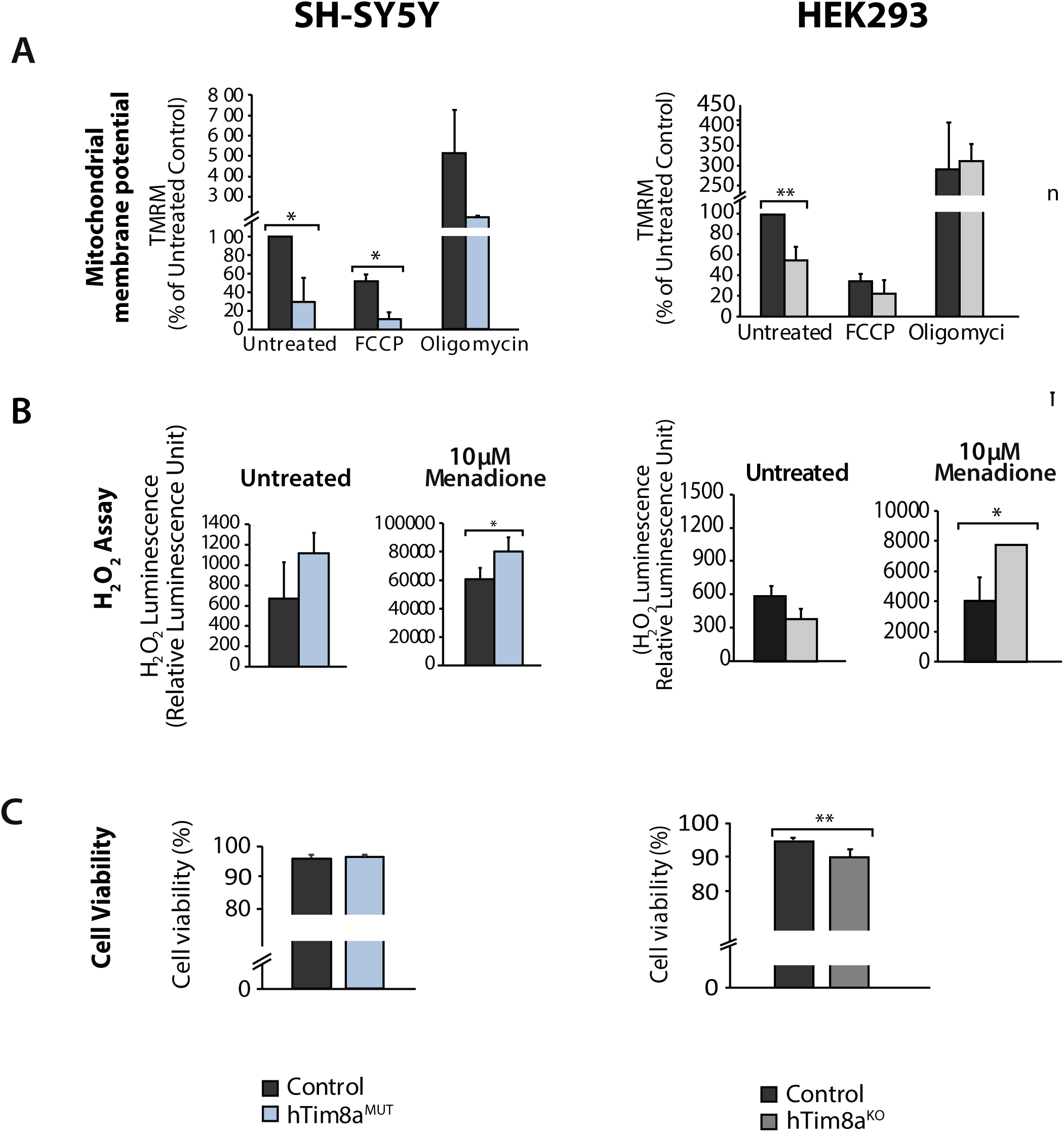
Mitochondrial dysfunction in hTim8a^KO^ HEK293 or hTim8a^MUT^ SH-SY5Y cells. (**A**) Mitochondrial membrane potential in hTim8a^MUT^ SH-SY5Y cells (left panel) or hTim8a^KO^ HEK293 cells (right panel) were quantified using TMRM. Cells were incubated with TMRM and, either: (i) DMSO (negative control); (ii) 20 μM FCCP to depolarise mitochondria; or (iii) 20 μM oligomycin to induce hyper-polarisation of the mitochondrial membrane potential. Data represents mean ± SEM (n=3). *, p< 0.05, **, p<0.01. (**B**) Reactive H_2_O_2_ species present in untreated or menadione-pretreated control, hTim8a^MUT^ SH-SY5Y cells (left panel) or hTim8a^KO^ HEK293 cells (right panel) were quantified using ROS-Glo™ H_2_O_2_ Assay. Data are shown as mean ± SD (n=3 for untreated, n=4 for menadione-treatment). *, p<0.05. (**C**) Cell viability of control and hTim8a^MUT^ SH-SY5Y cells (left panel) or hTim8a^KO^ HEK293 cells (right panel) were measured using Trypan Blue staining. Cells were seeded at the same confluency and 24 hours later were stained with trypan blue for cell counting. Viable cells were calculated as a percentage (1-(dead cells (stained blue)/ total number of cells) X 100 = % viable cells).

We looked at mitochondrial metabolism as an additional measure of mitochondrial and cellular health. Global changes in cellular and mitochondrial metabolism by metabolite profiling revealed a total of 200 metabolites by gas chromatography-mass spectrometry using a targeted MRM approach in both control and hTim8a^KO^ HEK293 cells. Fifty-nine of these metabolites were significantly altered (BH-adjusted p<0.05) in hTim8a^KO^ cells suggesting a distinct metabolomic footprint of these cells compared to control HEK293 (**Table 3**). Analysis of detected metabolites depicted as a heatmap with the top 40 most-significantly altered metabolites and as a pathway map (**Supplemental Figure 3B and 3C**), suggested significant changes in the central carbon metabolism of hTim8a^KO^ cells, indicated by decreases in the levels of almost all detected TCA cycle intermediates (including citrate, fumarate, malate), decreases in intermediates in lower glycolysis (DHAP, GAP, 2/3PGA, lactate) and a concomitant increase in intermediates in upper glycolysis and the pentose phosphate pathway (glucose, Glc6P, Rib5P, Ru5P). These results indicate that loss of hTim8a leads to specific defects in mitochondrial metabolism and alterations of non-mitochondrial pathways of central carbon metabolism in HEK293 cells.

Metabolite profiling indicated that the metabolic response of SH-SY5Y cells to loss of hTim8a, differed from that of hTim8a^KO^ HEK293 cells (**Figure 5A - 5D**). In particular, levels of many intermediates of the TCA cycle (citrate/isocitrate, fumarate, succinate) and interconnected pathways (glutamine) were significantly elevated, rather than reduced, in the hTim8a^MUT^ cells (**Figure 5A and 5C**). Exceptionally, both malate and aspartate were significantly reduced in hTim8a^MUT^ cells (**Figure 5A-5C**). The latter intermediates are involved in the malate-aspartate shuttle (**Figure 5B,** the most affected pathway in hTim8a^MUT^), which is responsible for transferring NADH reducing equivalents from the cytosol to the mitochondria, and is very active in brain tissues (McKenna et al., 2006). Another important feature that was apparent in hTim8a^MUT^ cells was the significant accumulation of epinephrine and striking changes in multiple metabolite intermediates of catecholamine pathway (metabolites/pathway coloured in pink in **Figure 5A and 5C**; phenylalanine and tyrosine metabolism in **Figure 5B**), a unique footprint observed in neuroblastoma cells lacking hTim8a but not in HEK293 cells. To further investigate whether loss of hTim8a in SH-SY5Y cells was associated with a defect in mitochondrial metabolic fluxes and the malate-aspartate shuttle, control and hTim8a^MUT^ SH-SY5Y cells were cultured in glucose-DMEM supplemented with [U-^13^C]-glutamine, or glutamine-DMEM supplemented with [U-^13^C]-glucose for 2 hours, and the time-dependent incorporation of ^13^C into the TCA cycle and related intermediates was quantified by GC-MS (**Figure 5D**). The catabolism of ^13^C-glucose and ^13^C-glutamine was highly compartmentalized in both cell lines, with ^13^C-glucose being catabolized via glycolysis to lactate, and ^13^C-glutamine being primarily catabolized in the TCA cycle (**Figure 5D**). Levels of ^13^C-enrichment in different intermediates in the two ^13^C-glucose labelled cell lines were similar, with the exception of glycerol-3-phosphate, which was more highly labelled in the hTim8a^MUT^ SH-SY5Y line (**Figure 5D**).

**Figure 5.**
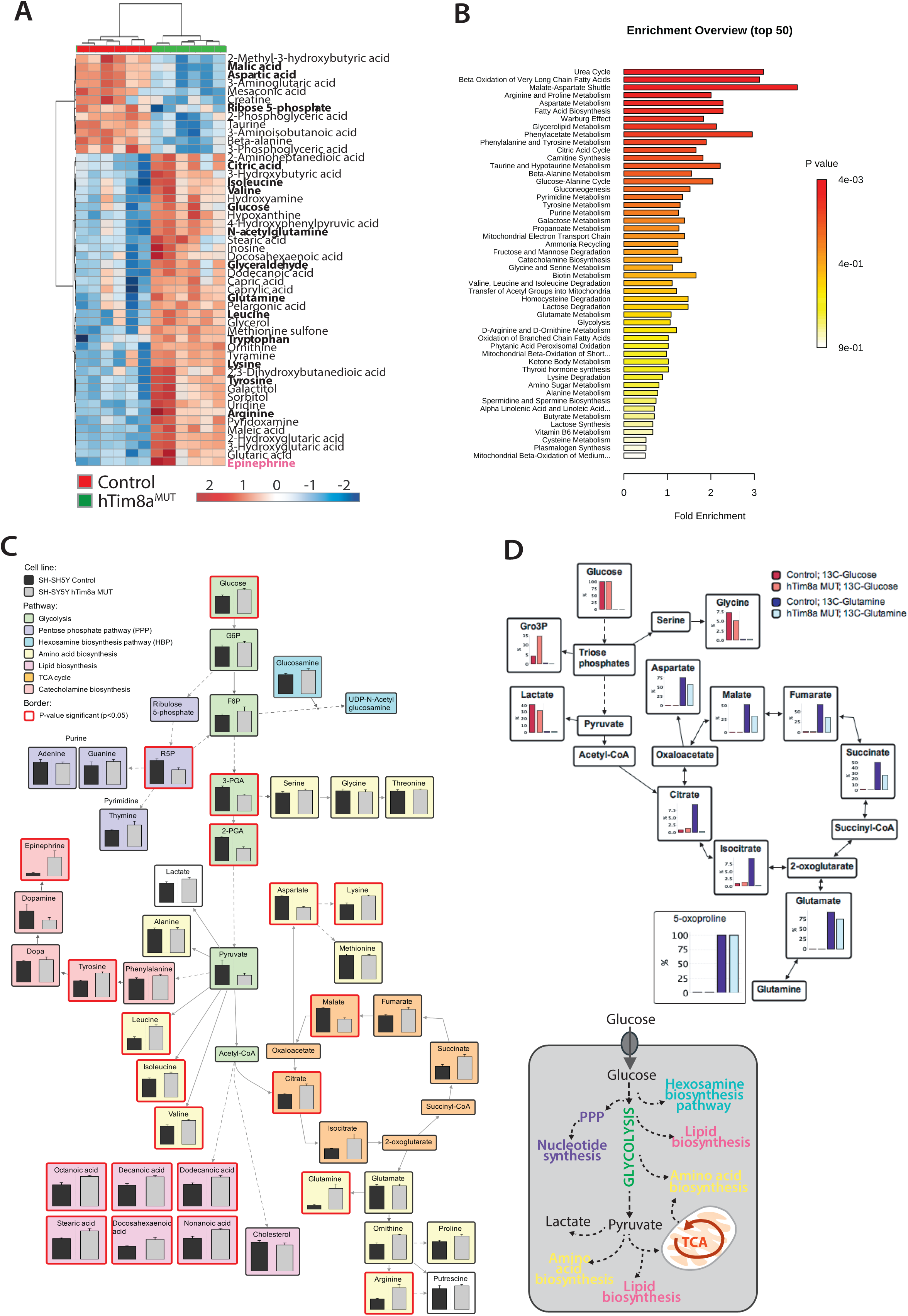
Cellular metabolism in hTim8a^MUT^ neuronal cells. (**A**) Relative abundance of significantly affected metabolites in hTim8a^MUT^ compared to control SH-SY5Y cells. Heatmap representing the top 50 most affected metabolites extracted from control and hTim8a^MUT^ cells (p< 0.05). Data represents values from 6 independent biological replicates of control and hTim8^MUT^ cells and fold changes are color-coded as indicated. See also **Table 3**. (**B**) Top 50 most affected cellular metabolic pathways in control and hTim8a^MUT^ cells were identified and represented as a clustered bar chart with its fold enrichment indicated. The p-value of each of the metabolic pathways are color-coded as indicated. (**C**) Relative abundances of intracellular metabolites were mapped onto metabolic networks using VANTED analysis tool. Each pathway is color-coded as indicated and p-value significant metabolites (p < 0.05) are boxed using a thicker, red-coloured border. See also **Table 4**. (**D**) Percentage of U-^13^C-glucose (indicated by red bars) or U-^13^C-glutamine (indicated by blue bars) labelled intracellular metabolites following a 2-hour incubation of control and hTim8a^MUT^ cells in ^13^C-glucose or ^13^C-glutamine labelled media. Data represents the mean of 3 independent experiments and are mapped onto their respective metabolic pathway. See also **Table 5.**

Significant differences were observed in the levels of ^13^C-enrichment in ^13^C-glutamine-fed control versus hTim8a^MUT^ SH-SY5Y cells. Glutamine feeds into the TCA cycle via the glutamine-glutamate-*α*-ketoglutarate pathway and can be used to drive mitochondrial respiration, and the synthesis of precursors for lipid and nucleotide biosynthesis, and neurotransmitters in neurons (Plaitakis et al., 2017). While levels of uptake of ^13^C-glutamine were not affected in the mutant line, levels of ^13^C-enrichment in intermediates in the oxidative TCA cycle, starting from *α*-ketoglutarate (as reflected in glutamate labelling) to malate/oxaloacetic acid were decreased by ∼30% compared to control cells (**Figure 5D**). Strikingly, ^13^C-enrichment in citric acid was almost completely repressed in the ^13^C-glutamine-fed hTim8a^MUT^ cells, indicating that most or all of the oxaloacetic produced in the mitochondria is exported to the cytosol as part of the malate-aspartate shuttle. Loss of hTim8a in SH-SY5Y cells therefore appears to lead to a decrease in the rate of utilization of glutamine, the major carbon source driving the mitochondrial TCA cycle and oxidative phosphorylation, leading to decreased production of malate and oxaloacetate and the pool of intermediates for the malate-aspartate shuttle. These analyses indicate that loss of hTim8a in SH-SY5Y cells leads to dysregulation of mitochondrial metabolism and redox balance, indicating a vital role in neuronal cell biology.

### Loss of hTim8a sensitises cells to intrinsic cell death

We set out to determine the cellular consequences of the observed mitochondrial dysfunction and changes to mitochondrial metabolism. The proteomic data (**Figure 1B**) of HEK293 cells lacking hTim8a highlighted changes to apoptotic regulators, including down-regulation in the levels of the Apoptosis Inducting Factor (AIF) and upregulation in the pro-apoptotic Bax. Changes to AIF and Bax were not evident in hTim9^MUT^ mitochondria, suggesting a specific cellular response due to the lack of hTim8a and a potential increased susceptibility to apoptosis. We confirmed changes to the levels of AIF and Bax by western blot and also uncovered a significant upregulation in the levels of cytochrome *c* (**Supplemental Figure 4A**). These changes were not off-target or clonal effects, as hTim8a^KO^ cells complemented with a tetracycline-inducible wild-type hTim8a (hTim8a^KO+WT^) restored the steady-state protein levels of Bax, cytochrome *c* and AIF (**Supplemental Figure 4B**). Alterations in the abundance of cytochrome *c* and Bax were not due to large changes in their gene expression as measured by qPCR (**Supplemental Figure 4A),** while the impact on AIF was due to transcriptional regulation. Treatment of control and hTim8a^KO^ cells with staurosporine, resulted in a faster release of cytochrome *c* from mitochondria in hTim8a^KO^ cells (**Supplemental Figure 4C**), suggesting these cells were indeed more sensitive to apoptotic induction compared to control cells (**Supplemental Figure 4D**). Given that the steady state viability of hTim8a^KO^ cells was not markedly compromised (**Figure 4C**), we propose that HEK293 cells lacking hTim8a are primed for cell death such that they can undergo a faster death upon cellular insult.

The implications of a primed for death state in neuronal cells would have greater implications than HEK293 cells as higher organisms including humans are limited in their ability to regenerate neuronal cells. We assessed if SH-SY5Y cells lacking hTim8 were adopting the same primed for death state. Mitochondria isolated from hTim8a^MUT^ cells had a significant increase in the level of cytochrome *c* relative to control cells (**Figure 6A**), but only modest changes in the steady-state levels of AIF and Bax were observed (**Figure 5B**). However, hTim8a^MUT^ SH-SY5Y cells were more vulnerable to staurosporine-induced apoptosis compared to control cells (**Figure 6B**). Apoptosis induction is usually accompanied by remodelling of the Bcl-2 family of proteins and caspase activation. To test if loss of hTim8a specifically leads to mitochondrial-mediated apoptotic cell death, we challenged the cells with ABT-737, a BH3 mimetic inhibitor of Bcl-xL, Bcl-2 and Bcl-w. Cell death and the rate of apoptosis were significantly increased in hTim8a^MUT^ cells and these effects were reversed when cells were pre-treated with a broad-spectrum caspase inhibitor QVD-OPh prior to addition of ABT-737 (**Figure 6C)**. In agreement with this, we observed increased Caspase-3 cleavage in hTim8a^MUT^ SH-SY5Y cells when treated with STS or ABT-737. Caspase-3 processing was completely inhibited in the presence of QVD-OPh (**Figure 6D, compare lanes 6 and 8 to 10 and 12**), consistent with the inhibition of apoptosis. Given that both apoptosis and ferroptosis are associated with ROS accumulation, we also examined whether loss of hTim8a increased vulnerability to this iron-dependent, lipid peroxidation-mediated form of regulated cell death (Dixon et al., 2012; Simon et al., 2000; Wu et al., 2018). Treatment with specific ferroptosis inducers, including: (i) Erastin, an SLC7A11 inhibitor; (ii) (1S,3R)-RSL3, which inhibits glutathione peroxidase 4 and (iii) buthionine sulfoximine (BSO), which depletes glutathione (GSH), did not enhance cell death of hTim8a^MUT^ SH-SY5Y cells (**Figure 6E**). We conclude that loss of hTim8a sensitises SH-SY5Y cells to Bcl-2-regulated and caspase-dependent, intrinsic cell death.

**Figure 6.**
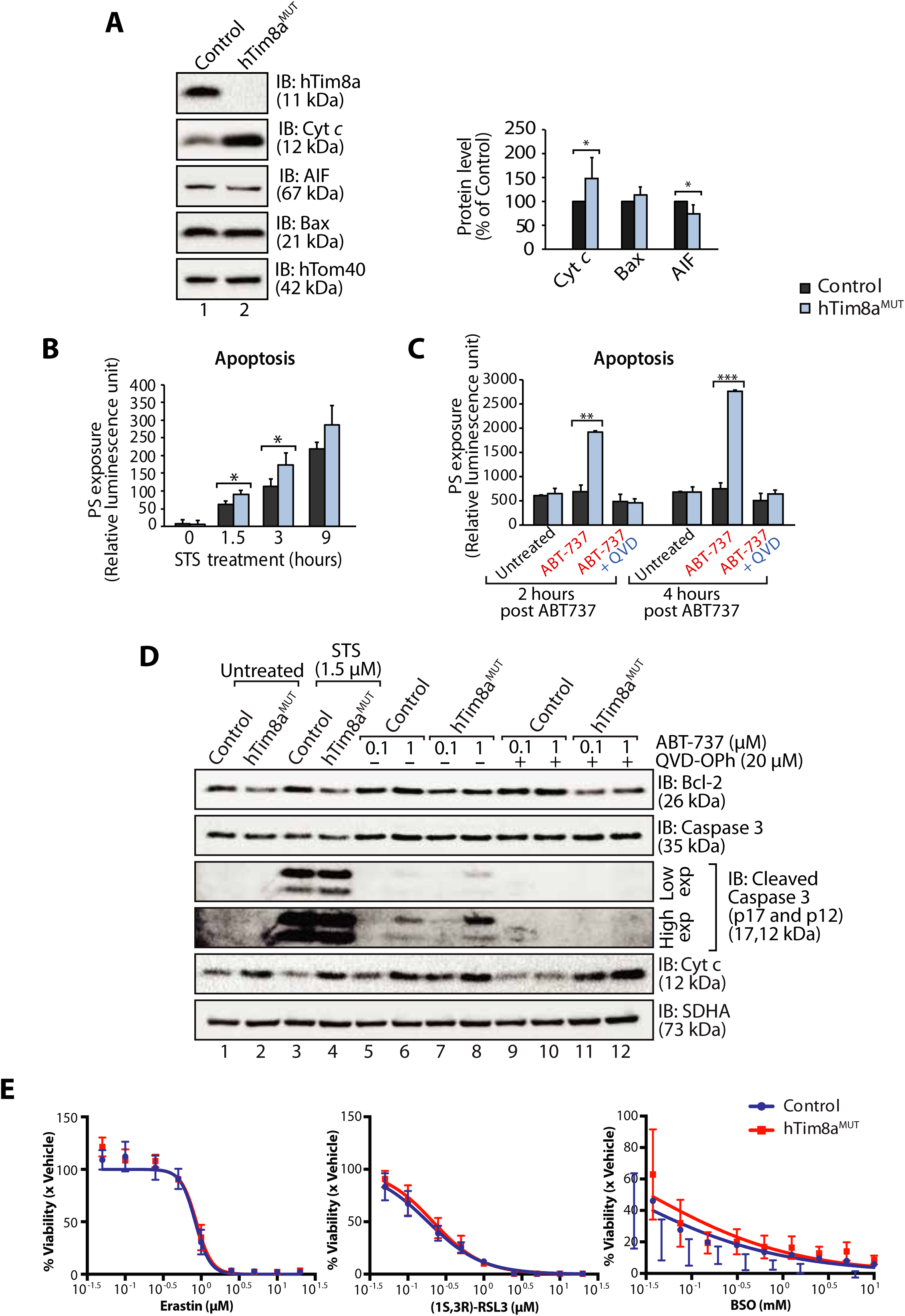
hTim8a^MUT^ cells are sensitive to apoptotic-cell death due to mitochondrial dysfunction. (**A**) Mitochondria isolated from control and hTim8a^MUT^ cells were analysed by SDS-PAGE and immunoblotting. Graph shows the relative levels of hTim23, Bax, Cyt *c* and AIF quantified and represented as mean ± SD (n=5). *, p<0.05. (**B**) Apoptotic sensitivity of control and hTim8a^MUT^ cells was measured following staurosporine (STS; 1.5 μM) treatment by assessing phosphatidylserine (PS) exposure (relative luminescence unit). n=4, mean ± SD; *, p<0.05. (**C**) Control and hTim8a^MUT^ cells was either: (i) left untreated, (ii) treated with ABT-737 (0.1 μM) or (iii) pretreated with QVD-OPh (20 μM) for 20 min prior to ABT-737 treatment, for 2 or 4 hours prior to measuring cellular apoptotic sensitivity by assessing phosphatidylserine (PS) exposure (relative luminescence unit). n=3, mean ± SD; **, p<0.01; ***, p<0.001. (**D**) Control and hTim8a^MUT^ cells were either (i) left untreated, (ii) treated with STS (1.5 μM) or (iii) treated with ABT-737 (0.1 or 1 μM) with or without preincubation with QVD-OPh. Cell lysates were harvested following these treatments for SDS-PAGE and immunoblot analyses using the indicated antibodies. (**E**) Control and hTim8a^MUT^ cells were treated with ferroptosis inducers at the indicated concentrations: Erastin (SLC7A11 inhibitor), (1S,3R)-RSL3 (GPX4 inhibitor) or BSO (γ-GCS inhibitor), for 72 hours to induce ferroptosis. The cellular viability following drug treatment was quantified using alamarBlue® assay and represented as mean ± SEM, n=4 biological replicates.

Given the Complex IV defects in hTim8a^MUT^ cells, we examined if the apoptotic vulnerability of cells lacking functional hTim8a was due to the observed oxidative stress. Challenging the cells with menadione alone increased the apoptotic vulnerability of hTim8a-deficient SH-SY5Y cells (**Figure 7A**), indicating that ROS accumulation and an impaired capacity to deal with ROS are contributing to the heightened apoptotic vulnerability in hTim8a^MUT^ cells. We wondered if attenuating the oxidative stress in hTim8a^MUT^ SH-SY5Y cells with anti-oxidants like Vitamin E (*α*-tocopherol; Vit E), an essential lipid soluble antioxidant that prevents lipid peroxidation in cellular membranes (Traber and Atkinson, 2007; Traber and Stevens, 2011), could lower their threshold to cell death induction. Although Vitamin E treatment did not restore the abundance of Complex IV subunits and assembly factors, MTCO1 and COX17 (**Figure 7B)**, or monomeric Complex IV (**Figure 7C**) in hTim8a^MUT^ mitochondria, it did did reduce the levels of cytochrome *c* (**Figure 7D**), which was accompanied by lowered sensitivity of these cells to apoptotic insult (**Figure 7E**). This was a Vitamin E specific response as treatment with Vitamin C, which functions as a water-soluble antioxidant (Traber and Stevens, 2011), did not rescue hTim8a^MUT^ cells for cell death vulnerability.

**Figure 7.**
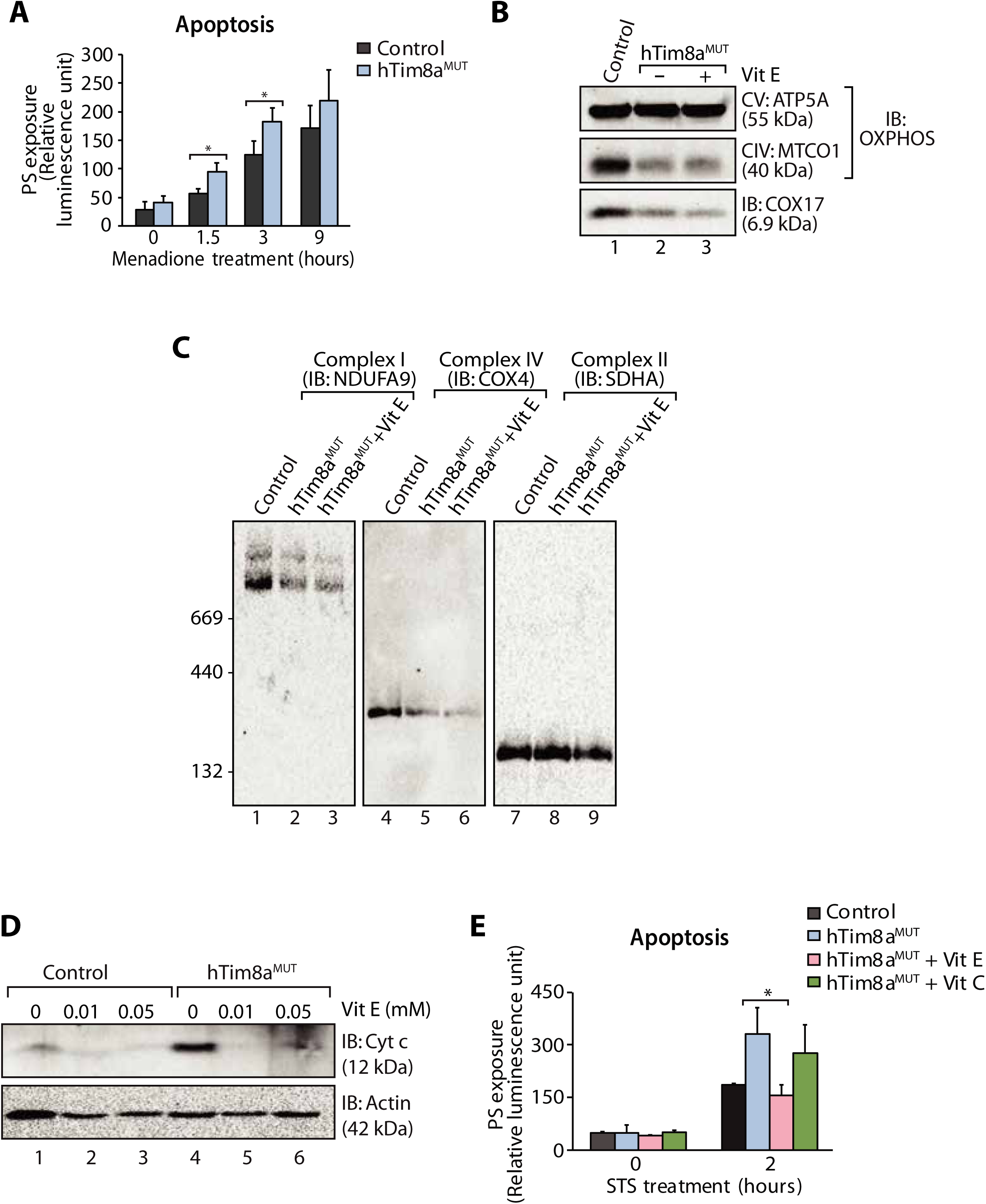
Elevated oxidative stress in hTim8a^MUT^ SH-SY5Y cells sensitises cells to apoptosis. (**A**) Apoptotic sensitivity of control and hTim8a^MUT^ cells was measured following menadione (10 μM) treatment by assessing phosphatidylserine (PS) exposure (relative luminescence unit). n=4 for control and n=3 for hTim8a^MUT^, mean ± SD; *, p<0.05. (**B**) Mitochondria were isolated from control and hTim8a^MUT^ cells following 24 hours of Vitamin E (Vit E, 0.01 mM) treatment prior to SDS-PAGE and immunoblotting for respiratory chain subunits. (**C**) Mitochondria were isolated from control, hTim8a^MUT^ treated with or without Vitamin E for 24 hours and solubilised in 0.5% TritonX-100-containing buffer before analysed using BN-PAGE/immunoblotting to detect respiratory chain complex stability. (**D**) Mitochondria were isolated from control and hTim8a^MUT^ cells following 24 hours of Vitamin E (Vit E, 0, 0.01 or 0.05 mM) treatment prior to SDS-PAGE and immunoblotting analyses. (**E**) Apoptotic sensitivity of control and hTim8a^MUT^ cells was measured following Vit C (0.2 mM) or Vit E (0.01 mM) treatment by assessing phosphatidylserine (PS) exposure (relative luminescence unit). n=3, mean ± SD; *, p<0.05.

## Discussion

This is the first study to comprehensively analyse the function of TIM chaperones across cell models and our findings highlight the functional specialisation of mitochondrial chaperones in different tissues, and open new opportunities for targeted nutritional therapies. Comparative analysis of neuronal-like and non-neuronal cells suggested that hTim8a facilitates the folding and/or transport of a number of substrate proteins, although this isoform was not required for assembly of the TIM22 complex or TIM22 protein import. In contrast, loss of hTim9 resulted in profound disruption of the TIM22-biogenesis pathway. Importantly, we also showed no impact to the biogenesis of hTim23 or the TIM23 complex, which has been suggested to be the underlying basis of MTS (Rothbauer et al., 2001).

Depletion of hTim8a in SH-SY5Y cells (hTim8a^MUT^) had a profound impact on the mitochondrial proteome, with a clear effect on Complex IV assembly. Complex IV is the terminal electron acceptor of mitochondrial respiratory chain and dysfunction of this complex is invariably associated with increased mitochondrial ROS and cellular toxicity. Complex IV deficiency causes a clinically heterogeneous variety of neuromuscular and non-neuromuscular disorders in childhood and adulthood and can result from either nuclear or mitochondrial mutations (Frazier et al., 2017). Complex IV is unique among the respiratory chain complexes as it has tissue-specific subunit isoforms, which are believed to have regulatory roles in adaptation of tissues to specific metabolic demands (Wallace and Fan, 2010). Six isoforms have been described for the nuclear-encoded COX subunits: three liver/heart-type specific pairs of isoforms (COX6A1/COX6A2, COX7A1/COX7A2, and COX8-1/COX8-2), the lung-specific isoform COX4-2, and two testes-specific isoforms, COX6B and COX8-3. The liver isoforms are found in tissues like brain that contain fewer mitochondria and we observed reduced levels of COX6A1 and COX7A2 in SH-SY5Y cells, but no changes to these subunits in the HEK293 background (hTim8a^KO^). COX17, which delivers Cu(I) to the assembly factors SCO1 and COX11 that in turn deliver Cu(I) to mt-DNA-encoded COX2, was one of few proteins reduced in all three hTim8a cell lines generated in this study: hTim8a^KO^ (HEK293), hTim8b^KO^ (HEK923) and hTim8a^MUT^ (SH-SY5Y) and hTim8b^KO^, hinting at a COX17-dependent process. Our data identifies a copper-independent but redox-sensitive interaction between hTim8a and COX17, which supports Complex IV biogenesis in SH-SY5Y cells. The presence of multiple Complex IV metallochaperones (SCO1), assembly factors (CMC1, COA4, COA6, COA7 and COX6B1) and core subunit (COX4l1) in the hTim8^FLAG^-COX17 cross-link suggests that hTim8a is binding to the COX17 pool participating in copper delivery to Complex IV, important for the maturation of Complex IV. In line with this, loss of functional COA6, which is also thought to be required for delivery of Cu(I) to COX2 (Pacheu-Grau et al., 2015; Stroud et al., 2015), results in a >1.5-fold increase in the levels of COX17, hTim8a, hTim8b and hTim13 (Stroud et al., 2015). Intriguingly, a similar pattern of Complex IV subunit turnover is observed in cells lacking COA6 to that observed in hTim8a^KO^ SH-SY5Y cells (Stroud et al., 2015), suggesting the proteins impact Complex IV at a similar stage of its assembly.

Cell lines lacking hTim8a display heightened vulnerability to apoptotic induction due to oxidative stress. We propose that cells lacking hTim8a are “primed” for cell death. The term “priming” is used to describe the proximity of cells to the apoptotic threshold (Ni Chonghaile et al., 2011) and the primed for death state is a continuum as the magnitude of BH3 proteins priming the mitochondrion can vary continuously until the anti-apoptotic reserve is overwhelmed and the cell commits to apoptosis (Certo et al., 2006). The apoptotic sensitivity of cells lacking hTim8a was selectively rescued by treatment with the lipid-soluble Vitamin E, but not the water-soluble Vitamin C. Vitamin E plays a major role in protecting membranes and nervous tissues from oxidative stress, where it scavenges peroxyl radical to maintain the integrity of long chain polyunsaturated fatty acids in cellular membranes (Traber and Atkinson, 2007; Traber and Stevens, 2011). Our results therefore suggest lipid peroxidation as a potential oxidative pathway modulated upon hTim8a loss, leading to oxidative stress-induced apoptosis. Importantly, MTS is characterized in its end-stage by widespread and severe neuronal cell death in the central nervous system (Hayes et al., 1998; Merchant et al., 2001; Tranebjaerg et al., 2001). In agreement with this, cells lacking hTim8a have significant defects in glucose catabolism and mitochondrial metabolism, both of which are evolutionarily linked to the regulation of cell death (King and Gottlieb, 2009). In SH-SY5Y cells, loss of hTim8a was associated with defects in glutamine catabolism and the operation of the malate-aspartate shunt, which may underlie global changes in redox balance and ROS production, further contributing to the acute severity of loss of hTim8a on neuronal function (Mergenthaler et al., 2013). Changes to the malate-aspartate shunt coincides with work showing that citrin and aralar1, calcium-responsive aspartate-glutamate carriers that function in the malate-aspartate shuttle, are substrates of the hTim8a-hTim13 complex (Roesch et al., 2004). Strikingly, loss of hTim8a in SH-SY5Y also skewed the levels of epinephrine (elevated levels) and dopamine (lowered levels), two principal neurotransmitters, which are synthesised by the catecholamine biosynthesis pathway, a reaction unique and essential to the central nervous system and brain (Kobayashi, 2001).

Our data demonstrates a cell-specific function for hTim8a in the assembly of Complex IV the SH-SY5Y cell model, an in vitro model for neuronal cell function. Previous work by Tranebjaerg et al. highlighted neuronal loss as a prominent feature in MTS patients (Tranebjaerg et al., 2001) and we now provide insight into the molecular features eliciting this cell loss and show cells lacking hTim8a are sensitised to an oxidative-stress induced intrinsic cell death. Importantly, our data highlights a strategy for early therapeutic intervention using Vitamin E for mitochondrial neuropathologies like MTS.

## Materials and Methods

### Cell lines and culturing, siRNA transfection, transient protein expression and stable cell line generation

Cell lines used in this work were HEK293T, Flp-In T-REx 293 (Thermo Fisher Scientific) and SH-SY5Y. Cells were cultured at 37 °C in Dulbecco’s modified Eagle’s medium (Gibco) containing 5% or 10% [v/v] fetal bovine serum (FBS; In vitro Technologies) and 0.01% penicillin-streptomycin [v/v] under an atmosphere of 5% CO_2_ and 95% air. Stable tetracycline inducible Flp-In T-REx HEK293 cell lines were generated using the T-Rex system (Thermo Fisher Scientific) as previously described (Kang et al., 2016). Briefly, cells plated overnight at 37 °C were transfected with pcDNA5/FRT/TO-(ORF) (plasmid encoding the ORF of interest) and pOG44 (encoding the Flp recombinase) at a 1:9 ratio (ng of DNA) using Lipofectamine 2000 prior to selection using 200 μg/ml of Hygromycin B (Thermo Fisher Scientific) for positive clones. To induce protein expression, cells were cultured in media supplemented with 1 μg/ml of tetracycline for the desired time.

### CRISPR/Cas9-gene editing and screening

pSpCas9(*gRNA*)-2A-GFP containing guide RNA against (i) *TIMM8A*, (ii) *TIMM9* or (iii) *TIMM8B* were transfected into Flp-In T-REx 293 or SH-SY5Y for generation of CRISPR/Cas9-edited cells (Ran et al., 2013). Cells were subjected to single cell sorting for GFP positive expression prior to expansion and screening for CRISPR/Cas9 gene editing using: (i) western blotting with antibody against specific protein; (ii) allele sequencing to identify specific mutations introduced into the gene of interest and (iii) complementation of CRISPR/Cas9-edited cell line with wild type protein to ensure no off-target effects.

Genomic sequencing of CRISPR/Cas9-edited clones revealed:

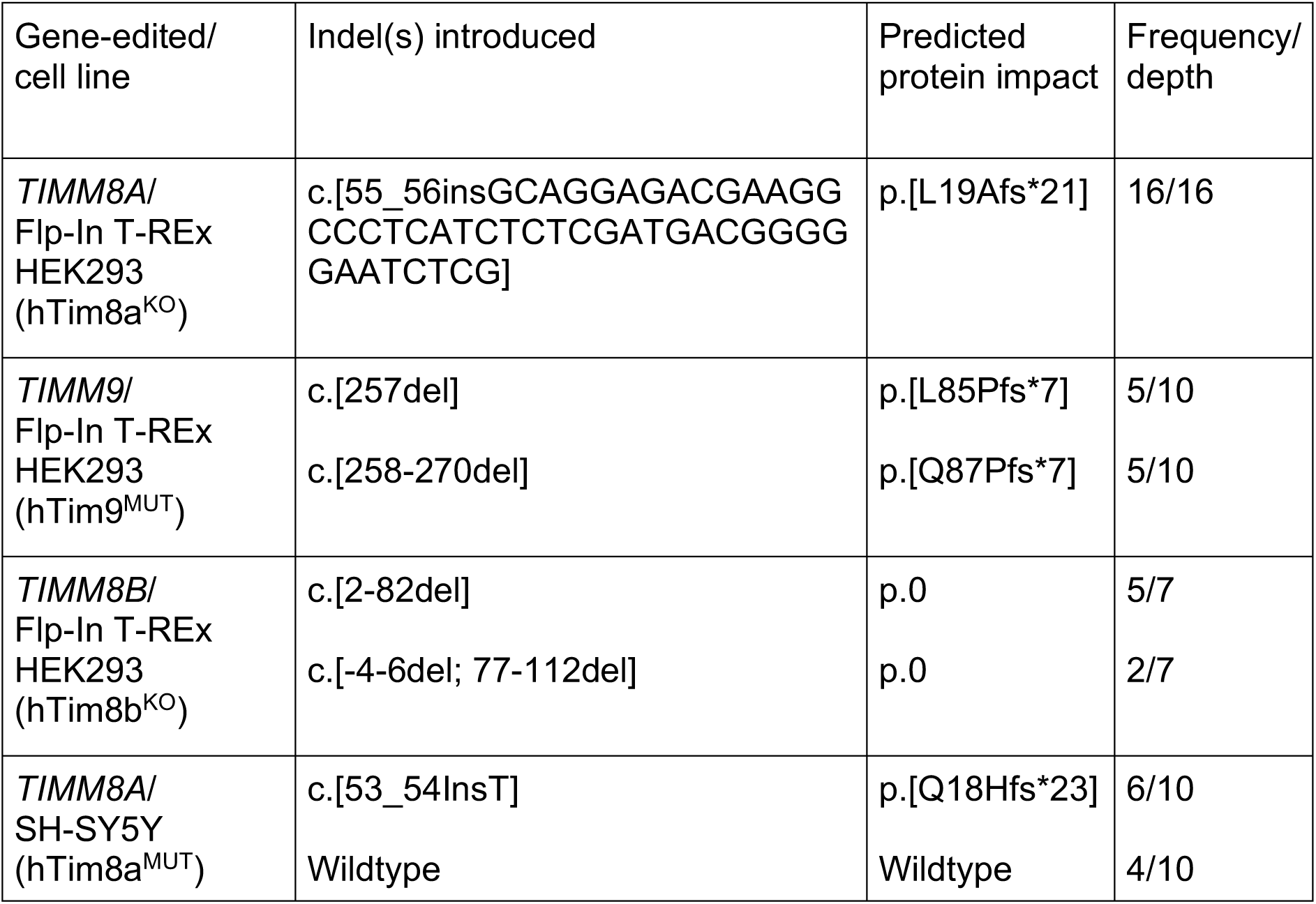

### Mitochondrial isolation and cell lysate preparation

Mitochondria were purified from tissue culture cells using differential centrifugation as previously described (Johnston et al., 2002; Kang et al., 2016; Kang et al., 2017). Cell pellets were homogenised in isolation buffer (20 mM HEPES-KOH, pH 7.4, 220 mM mannitol, 70 mM sucrose, 1mM EDTA, 0.5 mM PMSF, 2 mg/ml BSA) before centrifugation at 800 *g* to isolate cellular and nuclear debris. The supernatant was centrifuged at 12,000 *g* to obtain the mitochondria pellet. For preparation of whole cell lysate, cells were isolated and washed once in PBS prior to cell lysis in RIPA buffer (150 mM NaCl, 1 % Triton-X100, 0.5 % sodium deoxycholate, 0.1 % SDS, 50 mM Tris-Cl, pH 8.0). Protein concentrations were quantified using Pierce™ BCA protein assay kit (ThermoFisher Scientific) and desired amount of protein was TCA precipitated for SDS-PAGE analysis.

### Gel electrophoresis and immunoblot analysis

Tris-Tricine SDS-PAGE was carried out as described previously (Kang et al., 2016; Kang et al., 2017; Schagger and von Jagow, 1987). A 10-16% gradient was created using a gradient mixer. 16% and 10% acrylamide solutions (49.5% acrylamide: 1.5% bisacrylamide) were made up in tricine gel buffer (1 M Tris-Cl, pH 8.45, 0.1% [w/v] SDS) with 13% [v/v] glycerol added to the 16% mix. A stacking gel (4% acrylamide in tricine gel buffer) was overlayed onto the polymerised 10-16% gradient gel. Samples to be analysed were made up in SDS-PAGE loading dye (50 mM Tris-Cl, pH 6.8, 100 mM dithiothreitol, 2% [w/v] sodium dodecyl sulphate, 10% [w/v] glycerol, 0.1% [w/v] bromophenol blue). Electrophoresis was performed using Tris-tricine SDS-PAGE cathode buffer (0.1 M Tris, 0.1 M Tricine, pH 8.45, 0.1% SDS) and anode buffer (0.2 M Tris-Cl, pH 8.9).

Blue-Native-PAGE was performed as previously described (Ryan et al., 2001; Schagger, 1995). 4-16% or 4-13% gradient gels were typically utilised. Acrylamide solution (49.5% acrylamide: 1.5% bisacrylamide) consisting of 13% [v/v] glycerol in BN gel buffer (66 mM α-amino n-caproic acid, 50 mM Bis-Tris, pH 7.0) was used to generate a gradient gel, which was polymerised using TEMED and APS. A 4% [v/v] acrylamide solution in BN gel buffer was overlayed onto the gradient gel. Samples were solubilised in solubilisation buffer (20 mM Tris-Cl, pH 7.4, 50 mM NaCl, 0.1 mM EDTA and 10% [v/v] glycerol) with 1% [w/v] digitonin and incubated on ice for 30 min. 1:10 volume of 10X BN loading dye (5% [w/v] Coomassie blue G250 (MP Biomedicals, LLC), 500 mM α-amino n-caproic acid, 100 mM Bis-Tris pH 7.0) was added to the clarified supernatant prior to loading. Electrophoresis was carried out using BN cathode buffer (50 mM tricine, 15 mM Bis-Tris, pH 7.0, 0.02% [w/v] Coomassie blue G 250) and anode buffer (50 mM Bis-Tris, pH 7.0).

Following electrophoresis, gels were transferred onto 0.45 μm PVDF membranes using a semidry transfer apparatus before being subjected to immunoblot analysis using specific primary antibodies and secondary antibodies (Sigma). Protein detection was performed using ECL chemiluminescent reagent (GE Healthcare) on the ChemiDoc™ MP imaging machine (BioRad). Quantification of western blot signal was performed following the manufacturer’s instructions using the Image Lab software (BioRad). Details of primary antibodies used for western blotting analyses are described in the resource table.

### Crosslinking and immunoprecipitation

Mitochondria were resuspended in import buffer (20 mM HEPES-KOH (pH 7.4), 250 mM sucrose, 5 mM magnesium acetate and 80 mM magnesium acetate) supplemented with 0.5 mM ATP at 1 mg/ml and incubated with the amino-group specific homobifunctional and cleavable cross-linker, dithiobis(succinimidylpropionate) (0.2 mM) (DSP; Thermo Fisher Scientific) end-over-end for 1 hr at 4 °C. The reaction was quenched with 100 mM Tris-Cl, pH 7.4 for 30 min at 4 °C. Mitochondria were re-isolated by centrifugation at 16,000 g for 20 min at 4 °C and solubilised in lysis buffer (20 mM Tris-Cl, pH 7.4, 1 mM EDTA, 1% [w/v] SDS) with boiling at 95 °C for 5 min. Samples were clarified by centrifugation at 16,000 g for 5 min at RT before being diluted in 1% TritonX-100-containing buffer (1% [v/v] TritonX-100, 20 mM Tris-Cl, pH 7.4, 150 mM NaCl, 1 X complete protease inhibitor (Roche)). The sample was added to pre-equilibrated FLAG-resin and incubated end-over-end for 1 hr at 4 °C. The resin was isolated by centrifuging at 2,400 g for 3 min at 4 °C and unbound proteins were isolated. The resin and bound proteins were washed X4 in 1% TritonX-100 buffer before resuspending into 2X loading dye for SDS-PAGE and immunoblotting analyses.

### *In vitro* protein import and autoradiography

*In vitro* mRNA transcription and protein translation was performed using mMessage SP6 transcription kit (Ambion) and rabbit reticulocyte lysate system (Promega) respectively. *In vitro* translation was performed in the presence of ^35^S-methionine. Mitochondrial *in vitro* protein import assays were performed as described (Ryan et al., 2001). Radiolabeled proteins were incubated with isolated mitochondria resuspended at 1mg/ml in import buffer (20 mM HEPES-KOH (pH 7.4), 250 mM sucrose, 5 mM magnesium acetate and 80 mM magnesium acetate) supplemented with 10 mM sodium succinate, 1 mM 1,4-dithiothreitol and 5 mM ATP. Protein import was performed at 37 *°*C for the desired amount of time in the presence or absence of membrane potential. To dissipate the membrane potential, a final concentration of 10 μM of Carbonyl cyanide-p-trifluoromethoxyphenylhydrazone (FCCP; Sigma-Aldrich) was added to each import reaction. Import reaction was stopped at 4*°*C followed by incubation with 50 μg/ml of PK and subsequent treatment with 1 mM PMSF. Mitochondria were re-isolated and TCA precipitated for SDS-PAGE analysis or resuspended into digitonin-containing buffer for BN-PAGE analysis. Autoradiography was performed to visualise the radioactive signal using a Typhoon Phosphor Imager (GE Healthcare). Radioactive images were processed using Image J software.

### Quantitative mass spectrometry and data analysis

For mass spectrometry (MS) analysis, isolated mitochondrial protein pellet (200 μg) was solubilised in sodium deoxycholate (SDC) containing buffer (1% (w/v) SDC; 100 mM Tris-Cl, pH8.1; 40 mM chloroacetamide, 10 mM Tris(2-carboxyethy)phosphine (TCEP) followed by boiling at 99 *°*C for 15 min and sonication (Powersonic 603 Ultrasonic Cleaner, 40 KHz on high power) for 10 min at RT. Overnight digestion with 1:100 [w/w] of trypsin was performed at 37 *°*C. Samples were incubated with another 1:200 [w/w] trypsin for an additional 2 hours and tryptic peptides were extracted using 100 μl of ethyl acetate, 1% trifluoroacetic acid (TFA) and the entire volume was loaded onto 3M™Empore™ SDB-RPS stage tips (Kulak et al., 2014). The stage tips were washed twice using 200 μl of ethyl acetate, 1 % TFA followed by two washes with 200 μl of 0.2 % TFA. Peptides were eluted with 100 μl 80% acetonitrile, 5% ammonium hydroxide. All spins were performed at 1,500 × g at RT. Eluates were dried using a SpeedVac concentrator and peptides reconstituted in 15 μl of 2% acetonitrile (ACN), 0.1% TFA with sonication as above for 10 min. Samples were clarified at 16,000 *g* for 5 min at RT and soluble peptides transferred to autosampler vials.

Peptides were analysed by online nano-HPLC/electrospray ionization-MS/MS on Q Exactive Plus instruments connected to an Ultimate 3000 HPLC (Thermo-Fisher Scientific). For cell lines generated in the HEK293T background, peptides were loaded onto a trap column (Acclaim C18 PepMap nano Trap × 2 cm, 100 μm I.D, 5 μm particle size and 300 Å pore size; ThermoFisher Scientific) at 15 μL/min for 3 min before switching the pre-column in line with the analytical column (Acclaim RSLC C18 PepMap Acclaim RSLC nanocolumn 75 μm × 50 cm, PepMap100 C18, 3 μm particle size 100 Å pore size; ThermoFisher Scientific). The separation of peptides was performed at 250 nL/min using a 128 min non-linear ACN gradient of buffer A [0.1% (v/v) formic acid, 2% (v/v) ACN] and buffer B [0.1% (v/v) formic acid, 80% (v/v) ACN]. Data were collected in positive mode using Data Dependent Acquisition using m/z 375 – 1575 as MS scan range, HCD for MS/MS of the 12 most intense ions with z ≥ 2. Other instrument parameters were: MS1 scan at 70,000 resolution (at 200 m/z), MS maximum injection time 54 ms, AGC target 3E6, Normalized collision energy was at 27% energy, Isolation window of 1.8 m/z, MS/MS resolution 17,500, MS/MS AGC target of 2E5, MS/MS maximum injection time 100 ms, minimum intensity was set at 2E3 and dynamic exclusion was set to 15 sec. For the cell line generated in the SH-SH5Y background, peptides were loaded onto a trap column (PepMap C18 trap column 75 µm × 2 cm, 3 µm, particle size, 100 Å pore size; ThermoFisher Scientific) at 5 μL/min for 3 min before switching the pre-column in line with the analytical column (PepMap C18 analytical column 75 μm × 50 cm, 2 μm particle size, 100 Å pore size; ThermoFisher Scientific). The separation of peptides for was performed at 300 nL/min using a 185 min non-linear ACN gradient of buffer A [0.05% (v/v) TFA acid, 2% (v/v) ACN] and buffer B [0.05% (v/v) TFA, 80% (v/v) ACN]. Data were collected in positive mode using Data Dependent Acquisition using m/z 375 - 1400 as MS scan range, HCD for MS/MS of the 15 most intense ions with z ≥ 2. Other instrument parameters were: MS1 scan at 70,000 resolution (at 200 m/z), MS maximum injection time 50 ms, AGC target 3E6, Normalized collision energy was stepped at 25%, 30% and 35% energy, Isolation window of 1.6 m/z, MS/MS resolution 17,500, MS/MS AGC target of 5E4, MS/MS maximum injection time 50 ms, minimum intensity was set at 2E3 and dynamic exclusion was set to 30 sec.

Raw files were analysed using the MaxQuant platform (Tyanova et al., 2016a) version 1.6.1.0 searching against the Uniprot human database containing reviewed, canonical and isoform variants in FASTA format (May 2016) and a database containing common contaminants. Default search parameters for a label-free (LFQ) experiment were used with modifications. Briefly, “Match between runs” were enabled with default settings however .raw files from the two different instruments were given fraction numbers of 1 and 3 respectively to avoid spurious matching of MS1 peaks. Only unique peptides were used for quantification, using a LFQ minimum ratio count of 2. Using the Perseus platform (Tyanova et al., 2016b) version 1.6.1.1, proteins group LFQ intensities were Log2 transformed. Values listed as being ‘Only identified by site’, ‘Reverse’ or ‘Contaminants’ were removed from the dataset, as were identifications from <2 unique peptides. Annotations were imported from the IMPI Mitochondrial Proteome database (http://www.mrc-mbu.cam.ac.uk/impi) and ‘known mitochondrial’ proteins used for normalization of columns by row cluster. Experimental groups were assigned to each set of triplicates and a modified two-sided t-test based on permutation-based FDR statistics (Tyanova et al., 2016b) was performed. The negative logarithmic p-values were plotted against the differences between the Log_2_ means for the two groups. A significance threshold (FDR < 0.05, s0 = 0.5) was used for all experiments. Profile plots were generated as per Lake et al. (2017) (Lake et al., 2017). Briefly, mean and standard deviations were calculated for each experimental group in Perseus. Values from a group with a standard deviation > 0.3 were invalidated by conversion to “NaN” and rows were filtered to contain at least 2 valid values in both experimental groups. Tables were imported into Prism 7 software, following which a two-tailed ratio paired t-test was performed on the linearized Log2 LFQ Intensity mean values between the indicated groups.

### Metabolite extraction and GC-MS analysis

For steady-state metabolomics, cells were washed using phosphate-buffered saline (PBS) and frozen in liquid N_2_ directly in the tissue culture plate. Metabolite extraction was performed using methanol:chloroform (9:1; [v/v]) in the presence of 0.5 nmol ^13^C-Sorbitol and 5 nmol ^15^C_5_, ^15^N-valine as internal standards. Cells and supernatant were collected and centrifuged at 16,000 *g* at 4 *°*C for 5 min. Clarified supernatants were subjected to polar metabolite derivatization and analysis using a Shimadzu GC/MS-40 system (Best et al., 2018). Comprehensive targeted metabolite profiling data was processed using Shimadzu GCMS Browser software to generate a data matrix and was statistically analysed (Kang et al., 2017). Differences between the test and control samples were calculated using Student’s t-test to generate p-values that were further adjusted using Benjamini-Hochberg procedure to control for false-discovery rate (BH-adjusted p-value). To identify the relative flux of substrate into cells, ^13^C_6_-glucose or ^13^C_5_-glutamine labelling was performed as described (Kang et al., 2017; Kowalski et al., 2015). Briefly, cells were grown in glucose or glutamine-free media supplemented with 5 mM of ^13^C_6_-glucose or 4 mM ^13^C_5_-glutamine for 2 hours. Samples were then harvested for GC/MS and statistical analyses to determine the % of metabolite labelling as previously described. For metabolomic analyses, heatmaps and pathway mapping were generated using publicly available analysis tools: MetaboAnalyst 4.0 (Chong et al., 2018) or Vanted (Rohn et al., 2012).

### Gene expression with quantitative RT-PCR

Total RNA was extracted using NucleoSpin® RNA kit (Macherey-Negal), cDNA was synthesised from 1 μg of RNA using High-capacity cDNA Reverse Transcription kit (Applied Biosystems) and gene expression was determined by SYBR-green RT-PCR on the Lightcycler 480 (Roche). Gene expression was normalised to GAPDH and analysed using the *ΔΔ*C_t_ method. Primer sequences are detailed in the table below:

PCR primers for target genes:

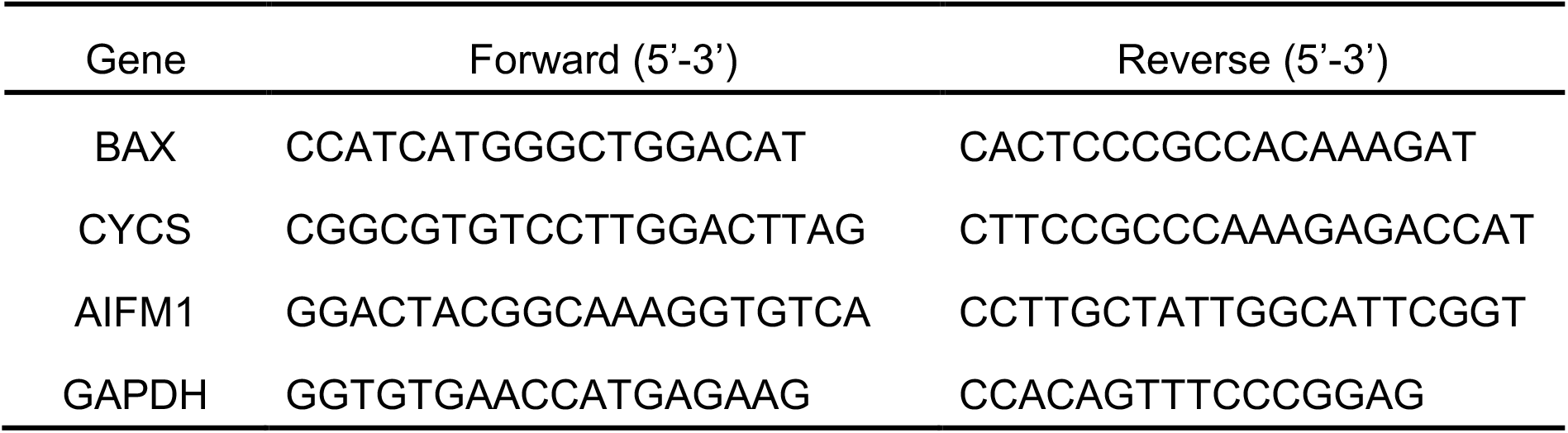

### Cellular viability measurements

Cell viability was assessed using trypan blue staining. 6 × 10^5^ cells were seeded and allowed to grow at 37 *°*C for 24 hours. Cells were harvested and stained with 0.4% trypan blue solution (ThermoFisher Scientific). Total number of cells and those stained blue were scored, and the number of viable cells was calculated by subtracting blue-stained (dead) cells from the total cells numbers. Conversely, % of viable cells was determined using the following formula: 1 - (Number of blue cells/Number of total cells) × 100%. Statistical significance of triplicate experiments was determined using two-tailed Student’s t-test.

The alamarBlue® assay was carried out as previously described (Liu et al., 2015). This assay quantifies the conversion of non-fluorescent resazurin to fluorescent by mitochondrial metabolic activity. For these experiments, SH-SY5Y cells were seeded in 96-well plates (1.5 × 10^4^ cells/well) and allowed to adhere overnight. The next day, these cells were treated with ferroptosis inducers: BSO (Sigma-Aldrich), Erastin or (1S,3R)-RSL3 (SelleckChem). Following 72 hours of drug treatment (at the indicated concentrations), 20 μL of 20% (v/v) alamarBlue® reagent (ThermoFisher Scientific) was added to each well without removing the pre-existing media. Cells were incubated for 2 h at 37°C, and fluorescence was measured using a FLUOstar OPTIMA microplate reader (BMG Labtech) at an excitation of 540 nm and an emission of 590 nm. The percentage of viable cells was calculated as follows 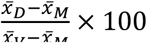, where 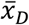: means fluorescence of drug-treated wells, 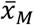: means fluorescence of media only wells, and 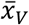: means fluorescence of vehicle-treated control wells.

### Apoptosis Assays

Apoptotic cell death was measured using RealTime-Glo™ Annexin V Apoptosis and Necrosis Assay (Promega), which measures phosphatidylserine (PS) exposure on the outer plasma membrane as a measure of apoptosis. 4 × 10^4^ cells were seeded onto a 96-well plate and incubated overnight at 37 *°*C. The next day, cells were (i) incubated with fresh media supplemented with or without staurosporine (1.5 μM; Sigma-Aldrich); (ii) pretreated with or without QVD-OPh caspase inhibitor (20 μM; Sigma Aldrich) for 20 minutes before incubated with ABT737 (1 μM); (iii) incubated with fresh media supplemented with or without menadione (10 μM; Sigma-Aldrich); or (iv) pretreated with Vitamin C or E (0.2 mM; 0.01 mM; Sigma) for 24 hours before staurosporine incubation. The detection reagent was added to a final concentration of 1X according to the manufacturer’s instructions. The relative fluorescence (necrosis) and luminescence (apoptosis) units were measured using FLUOstar OPTIMA microplate reader (BMG LABTECH) at the desired time interval following the different treatments. Differences in relative luminescence and fluorescence intensity between control and test samples were examined statistically using two-tailed Student’s t-test.

### Reactive Oxygen Species Measurement

Reactive oxygen species, H_2_O_2_ were measured using ROS-Glo™ H_2_O_2_ Assay according to the manufacturer’s instructions (Promega). 4 × 10^4^ cells were seeded onto a 96-well plate. Following 24 h of growth, H_2_O_2_ dilution buffer and substrate (provided by the manufacturer; to generate luciferin precursor) with or without 10 μM menadione (Sigma-Aldrich) was added to the cells and was incubated at 37 *°*C for 1 h incubation. ROS-Glo™ detection solution was added to the cells followed by a 20-min incubation at RT in the dark. Luminescence signals were measured using FLUOstar OPTIMA microplate reader (BMG LABTECH). Data generated from independent replicates of the experiment were analysed statistically using two-tailed Student’s t-test.

### Mitochondrial membrane potential measurement

Mitochondrial membrane potential was measured using Tetramethylrhodamine, methyl ester (TMRM; ThermoFisher Scientific) staining using a microplate reader. Briefly, cells were seeded at 4 × 10^4^ cells onto a 96-well plate and allowed to grow at 37 *°*C for 24 hours. Cells were treated with 75 nM TMRM stain, containing either DMSO (untreated), 20 μM FCCP or 20 μM Oligomycin for 30 min at 37 *°*C prior to incubation with the FCCP alone for another 30 min. The fluorescence signal of TMRM stain was detected at 590 nm with an excitation at 544 nm using FLUOstar OPTIMA microplate reader (BMG LABTECH).

### Citrate Synthase Assay

To measure citrate synthase (CS) activity, cells grown in triplicate flasks were homogenized and centrifuged at 600 *g*. CS activity was determined in the post-600 *g* supernatants as described earlier (Frazier and Thorburn, 2012) using spectrophotometry. This assay measures the rate of free sulfhydryl group production using the thiol reagent 5,5-ditio-bis-(2-nitronenzoic acid) which reacts with the sulfhydryl group to form a free thionitrobenzoate anion. Briefly, CS buffer was loaded into the cuvette (50 mM KPi, pH 7.4, 0.1 mM 5,5’-dithio-bis-(2-nitrobenzoic acid); equilibrated to 30 °C) prior to the additional of the sample and Acetyl-CoA to a final concentration of 0.1 mM. The reaction was initiated with the addition of oxaloacetic acid (0.1 mM final concentration). Absorbance of thionitrobenzoate anion produced was measured at 412 nm using spectrophotometer and the linear reaction rate was obtained for about 3 mins. Spectrophotometric trace was then analysed and linear regions was selected for calculating initial rates of enzyme activity in nmol/min/mg protein.

### Antibodies

| Antibody | Company | Catalogue Number |
| --- | --- | --- |
| Rabbit polyclonal AGK | Atlas Antibodies | HPA020959 |
| Mouse monoclonal hTim9 | Abcam | ab57089 |
| Mouse monoclonal SDHA | Abcam | ab14715 |
| Rabbit polyclonal ANT3 | Abcam | ab154007 |
| Rabbit polyclonal<br>Glutamate carrier 1 (GC1) | Abcam | ab137614 |
| Mouse monoclonal<br>OXPHOS | Abcam | ab110413 |
| Mouse monoclonal<br>ATP5alpha | Abcam | ab14748 |
| Rabbit polyclonal hTom70 | Abcam | ab83841 |
| Mouse monoclonal<br>Cytochrome c | BD Biosciences | 556433 |
| Mouse monoclonal<br>hTim23 | BD Biosciences | 611223 |
| Rabbit polyclonal COX4 | Cell Signaling<br>Technology | 4850 |
| Rabbit monoclonal AIF | Cell Signaling<br>Technology | 5318 |
| Rabbit polyclonal<br>Caspase-3 | Cell Signaling<br>Technology | 9662 |
| Rabbit polyclonal hTim8a | Proteintech | 11179-1-AP |
| Rabbit polyclonal hTim13 | Proteintech | 11973-1-AP |
| Rabbit polyclonal hTim44 | Proteintech | 13859-1-AP |
| Rabbit polyclonal COX17 | Sigma Aldrich | HPA048158 |
| Mouse monoclonal FLAG | Sigma-Aldrich | F1804 |
| Rabbit polyclonal hTim22 | Sigma-Aldrich | T8954 |
| Rabbit polyclonal Tim29 | Sigma-Aldrich | HPA041858 |
| Mouse monoclonal actin | Sigma-Aldrich | A2228 |
| Mouse monoclonal Bcl-2 | ThermoFisher Scientific | 13-8800 (Bcl-2-100) |
| Mouse monoclonal<br>hTom22 | SantaCruz | Sc-58308 |
| Rabbit polyclonal Sam50 | N/A | Mike Ryan, Monash University |
| Rabbit polyclonal NDUFAF2 | N/A | Mike Ryan, Monash University |
| Rabbit polyclonal hTom40 | N/A | Mike Ryan, Monash University |
| Rabbit polyclonal Mfn2 | N/A | Mike Ryan, Monash University |
| Rabbit polyclonal NDUFV2 | N/A | Mike Ryan, Monash University |
| Rabbit polyclonal NDUFA9 | N/A | Mike Ryan, Monash University |
| Rat monoclonal BAX | N/A | Grant Dewson, Walter and Eliza Hall Institute |

## Acknowledgments

YK is supported by Melbourne International Fee Remission Scholarship (MIFRS) and Melbourne International Research Scholarship (MIRS). DS is supported by a Research Fellowship from the Mito Foundation. We acknowledge funding from the Australian Research Council (DP170101249 to DS), NHMRC Project Grants (1125390, 1107094 to MTR., DRT., and DAS; 1140906 to DAS and MTR) and Fellowships (1140851 to DAS; 1022896 to DRT; and 1059530 to MJM), Victorian Government Department of Health and Human Services acting through the Victorian Cancer Agency (MCRF16002 to NJC), the Mito Foundation and the Victorian Government’s Operational Infrastructure Support Program. We thank Simone Tregoning for research support. We thank the Bio21 Mass Spectrometry and Proteomics Facility and the Monash University Biomedical Proteomics Facility for the provision of instrumentation, training, and technical support. We acknowledge the use of the Biological Optical Microscopy Platform at the University of Melbourne. We thank A/Prof. Grant Dewson for discussion and reagents.

## Declaration of Interests

The authors declare no competing interests.

---

**Supplemental Figure 1:**
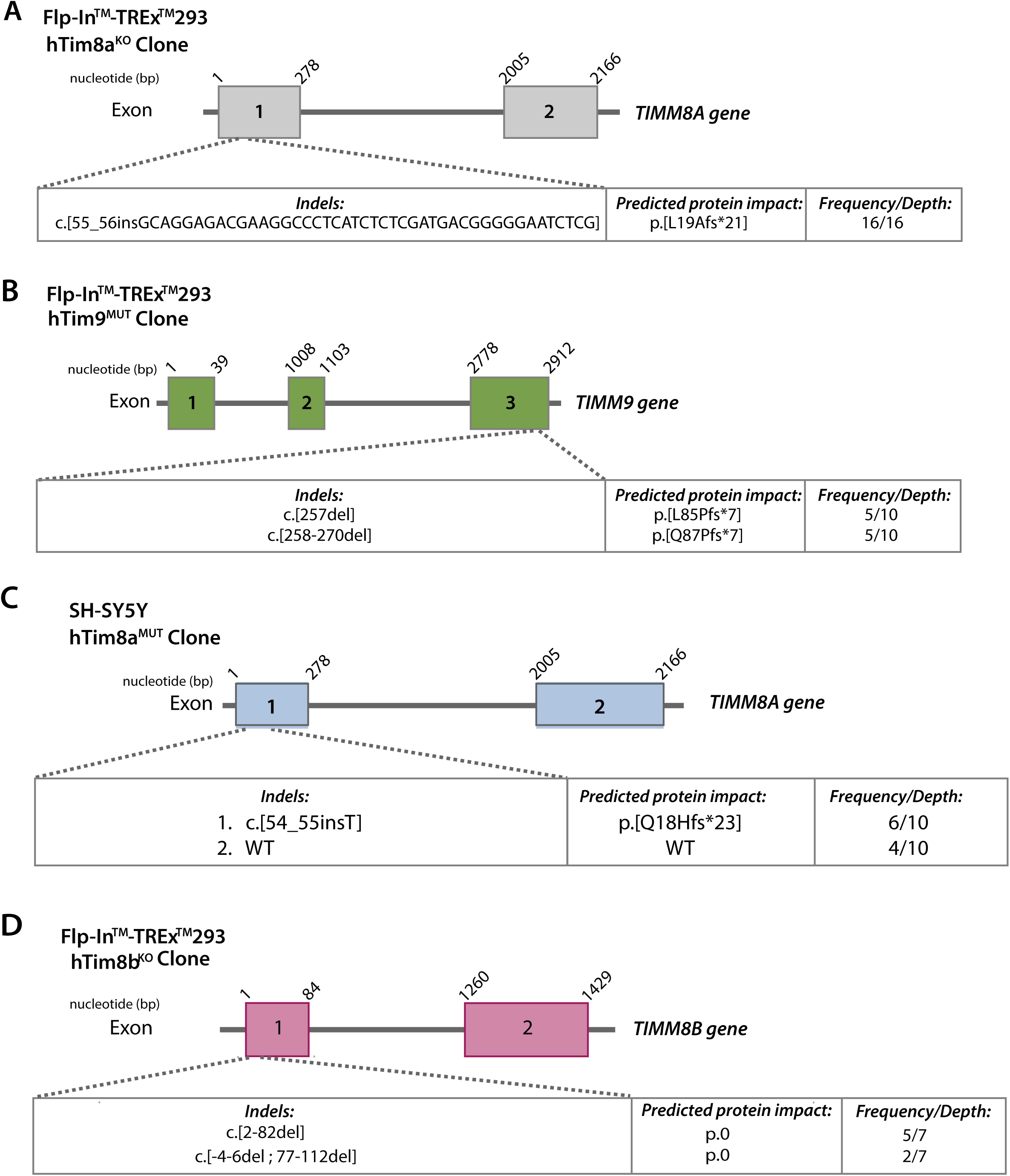
**Generation of CRISPR/Cas9 genome-edited cell lines.** (**A-D**) Schematic representation of the CRISPR/Cas9 editing system used to generate (A) hTim8a^KO^ (knockout), (B) hTim9^MUT^ cells in Flp-In™ T-Rex™ HEK293 cells, (C) hTim8a^MUT^ (mutant) in SH-SY5Y cells or (D) hTim8b^KO^ cells in Flp-In™ T-Rex™ HEK293 cells.

**Supplemental Figure 2:**
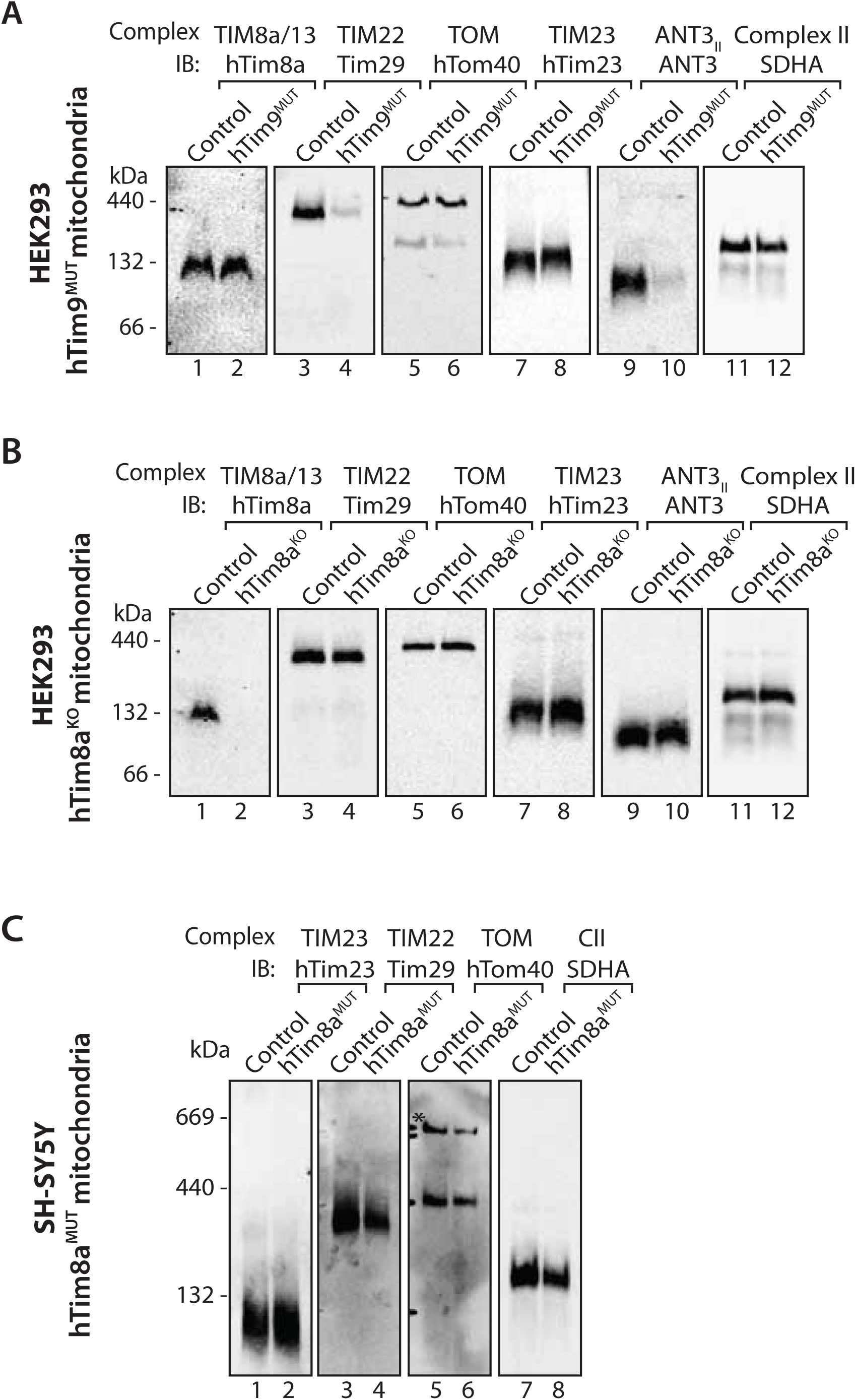
**Loss of hTim8a is associated with destabilisation of the hTim8a/hTim13 complex but no effect on TIM complex, Related to Figure 1.** (**A-C**) Mitochondria isolated from control and (**A**) hTim9^MUT^ HEK293 cells, (**B**) hTim8a^KO^ HEK293 cells or (**C**) hTim8a^MUT^ SH-SY5Y cells, were lysed in 1% digitonin prior to BN-PAGE analysis and western blotted using the indicated antibodies.

**Supplemental Figure 3:**
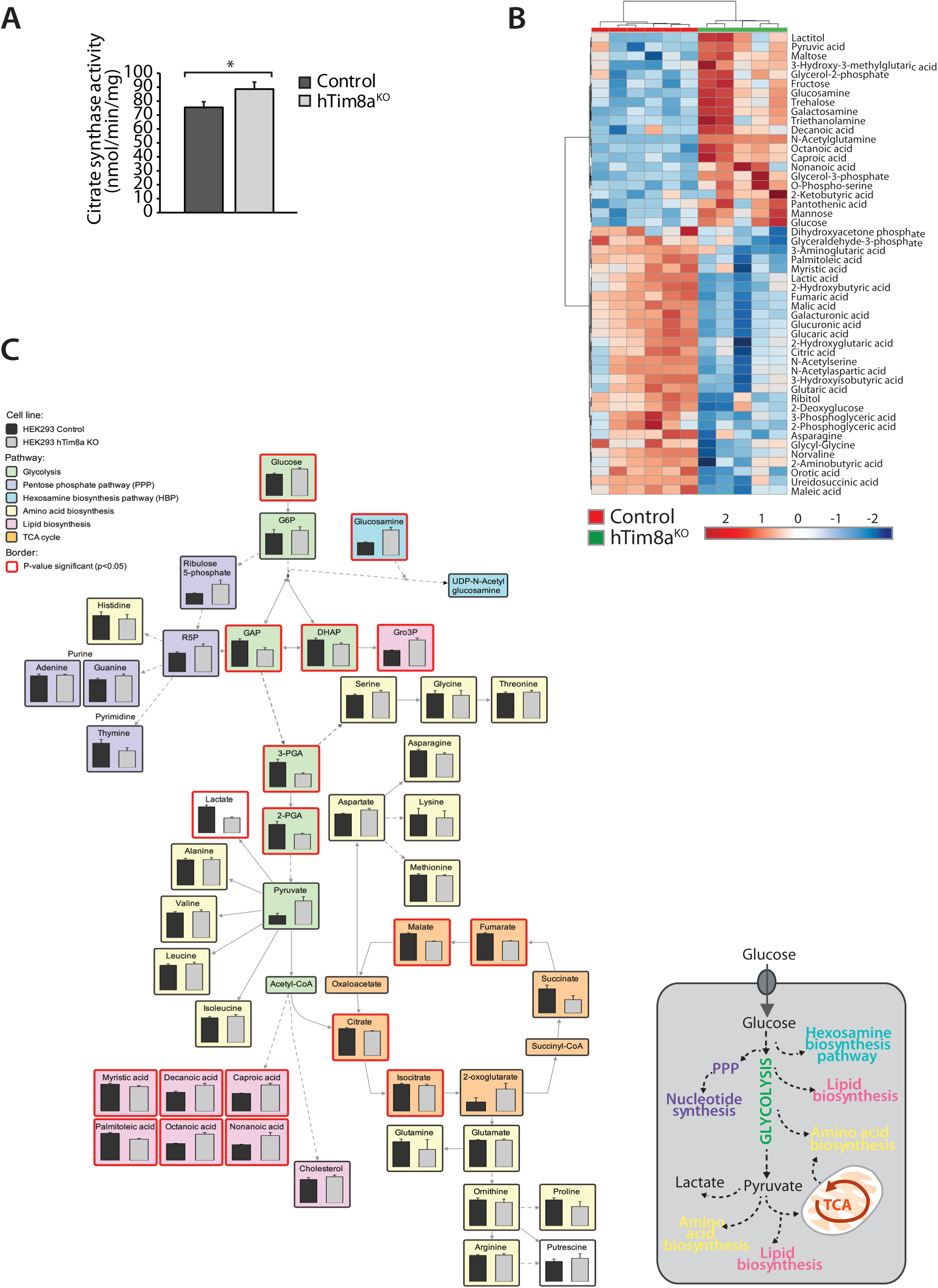
**Metabolic dysfunctions in hTim8a^KO^ HEK293 cells,** (**A**) The activity of mitochondrial marker enzyme, citrate synthase in control and hTim8a^KO^ cells was measured by spectroscopy in whole cell lysates from control and hTim8a^KO^ cells. Data is shown as mean ± SD (n=3). *, p< 0.05. (**B**) Hierarchical clustering of relative abundance of the top 40 significantly-affected intracellular metabolites depicted as a heat map (p-value < 0.05) in control and hTim8a^KO^ HEK293 cells. See also **Table 3**. (**C**) Relative abundance of intracellular metabolites in control and hTim8a^KO^ cells was mapped onto networks using VANTED analysis tool. Pathways are color-coded as indicated and significant metabolites (p < 0.05) are boxed using a thicker, red-coloured border.

**Supplemental Figure 4:**
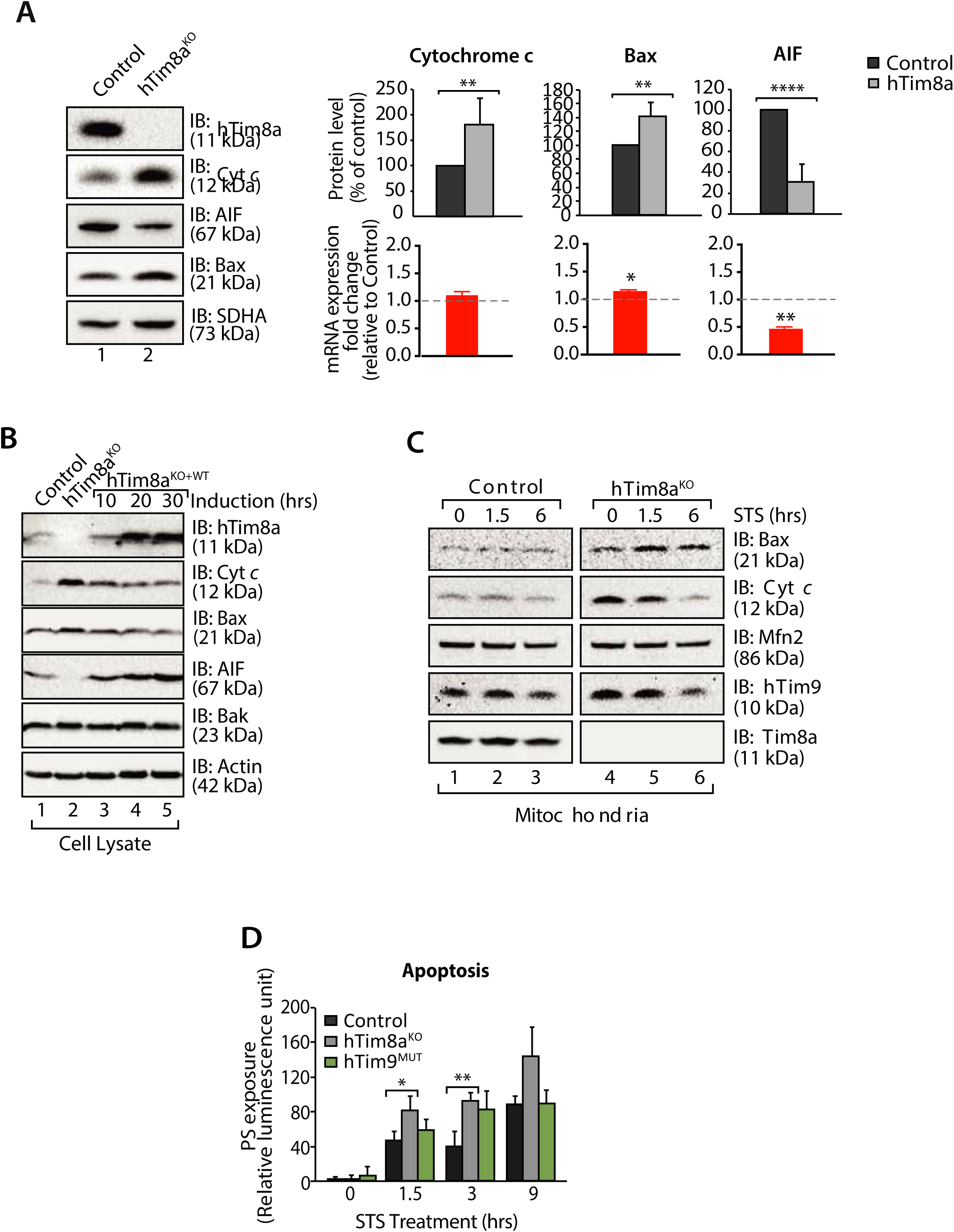
**hTim8a^KO^ HEK293 cells are more sensitive to cell death induction** (**A**) Mitochondrial lysates were analysed by SDS-PAGE and western blotting. Relative protein levels of Bax, cytochrome *c* (Cyt *c*) and apoptosis-inducing factor (AIF) were quantified and tabulated as mean ± SD (n=3). Relative fold change of mRNA expression of Bax, Cyt *c* and AIF in hTim8a^KO^ compared to control HEK293 were quantified and tabulated as mean ± SEM (n=3) (**B**) Cell lysates from control, hTim8a^KO^ and hTim8a^KO^ cells re-expressing hTim8a (hTim8a^KO+WT^) were analysed using SDS-PAGE and immunoblotting with the indicated antibodies. (**C**) Control and hTim8a^KO^ cells were treated with staurosporine (STS; 1.5 μM) for 0, 1.5 or 6 hours prior to mitochondrial isolation and analysis by SDS-PAGE. (**D**) Control, hTim8a^KO^ and hTim9^MUT^ cells were treated with staurosporine (STS; 1.5 μM) for the indicated time. Rate of apoptosis was calculated by measuring phosphatidylserine (PS) exposure to the outer leaflet of plasma membrane (relative luminescence unit). n=3, mean ± SD; *, p<0.05; **, p<0.01; ****, p<0.0001.

